# *APOE4* exacerbates glucocorticoid stress hormone-induced tau pathology via mitochondrial dysfunction

**DOI:** 10.1101/2025.02.03.636364

**Authors:** Qing Yu, Fang Du, Jeffrey Goodman, Clarissa L. Waites

**Author notes:** These authors contributed equally to this work (Qing Yu, Fang Du). Correspondence to: Clarissa Waites, Ph.D., 650 W. 168^th^ St., Black Building 1210B, New York, NY 10032, USA.

## Abstract

*APOE4* is the leading genetic risk factor for Alzheimer’s disease, and chronic stress is a leading environmental risk factor. Studies suggest that *APOE4* confers vulnerability to the behavioral and neuropathological effects of chronic stress, representing a potential mechanism by which this genetic variant accelerates Alzheimer’s onset and progression. Whether and how *APOE4*-mediated stress vulnerability manifests in neurons of the hippocampus, a brain region particularly susceptible to stress and Alzheimer’s pathology, remains unexplored.

Using a combination of *in vivo* and *in vitro* experiments in humanized *APOE4* and *APOE3* knockin mice and primary hippocampal neurons from these animals, we investigate whether and how *APOE4* confers sensitivity to glucocorticoids, the main stress hormones.

We find that a major hallmark of stress/glucocorticoid-induced brain damage, tau pathology (i.e., tau accumulation, hyperphosphorylation, and spreading) is exacerbated in *APOE4* versus *APOE3* mice. Moreover, *APOE4* animals exhibit underlying mitochondrial dysfunction and enhanced glucocorticoid receptor activation in the hippocampus, factors that likely contribute to tau pathogenesis in both the presence and absence of stress/glucocorticoids. Supporting this concept, we show that opening of the mitochondrial permeability transition pore drives mitochondrial dysfunction and tau pathology in *APOE4* mice, and that pharmacological inhibition of pore opening is protective against ApoE4-mediated mitochondrial damage, tau phosphorylation and spreading, and downstream hippocampal synapse loss. These findings shed light on the mechanisms of stress vulnerability in *APOE4* carriers and identify the mitochondrial permeability transition pore as a potential therapeutic target for ameliorating Alzheimer’s pathogenesis in this population.

## Introduction

Alzheimer’s disease is caused by a complex interplay of genetic and environmental factors. The strongest genetic risk factor for late-onset Alzheimer’s is *APOE4*, encoding the E4 variant of the lipid transporter apolipoprotein E (ApoE)^1^. *APOE4* is carried by ∼14% of the world’s population and increases Alzheimer’s disease risk by 4-fold (for heterozygotes) or 12-fold (for homozygotes) compared to the risk-neutral *APOE3* variant^1^. While environmental risk factors are harder to pinpoint, multiple studies indicate that chronic stress and elevated levels of glucocorticoids (GCs), the main stress hormones, increase Alzheimer’s risk and hasten disease progression^2-4^. Indeed, prolonged psychological stress during mid-life, high work-related stress, and high ‘distress proneness’ during middle age are all reported to significantly elevate Alzheimer’s risk, while high circulating levels of glucocorticoids (GCs), the major stress hormones, are associated with faster cognitive decline in Alzheimer’s subjects^2,3,5-7^. Stress-related neuropsychiatric disorders, particularly depression, are also linked to higher Alzheimer’s and dementia risk^8,9^. Interestingly, *APOE4* carriers exhibit a higher incidence of late-life depression, anxiety, and cognitive impairment compared to carriers of the other *APOE* isoforms^10-13^, and these conditions are exacerbated to a greater extent by environmental stressors^10,14^. These findings suggest that *APOE4* confers stress vulnerability, representing a potential mechanism by which this gene accelerates Alzheimer’s disease onset and progression. Whether and how such stress vulnerability manifests at the cellular level to promote disease pathogenesis remains unclear.

Two prominent drivers of Alzheimer’s disease are mitochondrial dysfunction and tau pathogenesis (i.e., hyperphosphorylation, oligomerization, and trans-cellular spreading of tau)^4,15,16^. We previously showed that high GC levels strongly induce both of these features in the murine hippocampus and that they are linked, with GC-induced mitochondrial dysfunction promoting tau pathology^17^. ApoE4 expression also promotes these features. Indeed, the brains of *APOE4* carriers (humans and targeted replacement *APOE4* knockin mice) exhibit altered mitochondrial structure and function, including impaired mitochondrial fission and mitophagy, elevated levels of reactive oxygen species (ROS) and oxidative stress markers, and decreased mitochondrial membrane potential and respiration^18-21^. Interestingly, *APOE4* mice also exhibit age- and stress-induced depressive behaviors that are rescued by supplementation with ATP^20^, suggesting that disruption of mitochondrial ATP production contributes to stress vulnerability in *APOE4* carriers. Moreover, *APOE4* exacerbates tau hyperphosphorylation and tau-mediated neuronal loss and brain atrophy in the PS19 tauopathy mouse model^22^, and *APOE4* deletion from brain cells of these animals significantly mitigates these phenotypes^23,24^. Collectively, these studies indicate that ApoE4 expression disrupts mitochondrial function and promotes tau pathology, although the link between these two processes in *APOE4* carriers remains unclear.

In the current study, we examine the effects of GCs on tau pathogenesis and mitochondrial function in *APOE4* vs *APOE3* humanized knockin mice. We find that GC-induced tau pathology, including tau accumulation, phosphorylation, and spreading, is exacerbated in *APOE4* vs. *APOE3* carriers. Middle-aged *APOE4* animals also exhibit oxidative stress, mitochondrial dysfunction, and enhanced GR activation in the hippocampus under control conditions, suggesting that these underlying factors contribute to tau pathogenesis in the presence and absence of GCs. Further, our findings implicate the mitochondrial permeability transition pore (mPTP) in these processes, and show that mPTP inhibition is protective against both mitochondrial damage and tau pathology in *APOE4* mice. These findings shed light on the mechanisms of stress vulnerability and Alzheimer’s disease pathophysiology in carriers of the *APOE4* gene.

## Materials and Methods

### Mice

APOE3 (strain #029018), APOE4 (strain #027894) and PS19 (strain #008169) mice were obtained from The Jackson Laboratory. All animal studies were carried out with the approval of the Columbia Institutional Animal Care and Use Committee (IACUC) in accordance with the National Institutes of Health guidelines for animal care. Offspring of transgenic (Tg) mice: APOE3(*+/+*)(E3), APOE4(*+/+*)(E4), PS19(*+/-*)/APOE3(*+/+*)(TE3), and PS19(*+/-*)/ APOE4(*+/+*)(TE4) mice were identified by PCR using primers for each specific transgene. Mice were administered the following drugs: DEX [D2915, Sigma; 5 mg/kg per day by intraperitoneal (IP) injection, dissolved in PBS], mito-apocynin [MitoAPO, HY-135869, MedChemExpress; 3 mg/kg per day by IP injection, dissolved in 15% polyethylene glycol (PEG400) with PBS]. 15 days DEX/mito-apocynin injection for 9-10 months/15-16 months APOE3/APOE4 (E3/E4) mice, 10 weeks mito-apocynin injection for 1-1.5 months PS19/APOE4 (TE4) mice. Control animals received daily intraperitoneal injections of PBS (dex vehicle) or 15% PEG400 in PBS (mito-apocynin vehicle). Both male and female mice were used in the experiments.

### Evaluation of serum corticosterone levels

Endogenous corticosterone serum levels were measured using the Corticosterone Parameter Assay kit (R&D Systems, KGE009) as described previously^25^. The wavelength for measurement was 450 nm and the correction wavelength was 570 nm.

### Primary hippocampal culture

Primary neurons were prepared from postnatal (P0-1) APOE3 or APOE4 hippocampi, and maintained in 24-well plates with Neurobasal medium supplemented with B27, 600 μM L-glutamine, and antibiotic-antimycotic, as described previously^26^. At 11-12 days *in vitro* (DIV), hippocampal neurons were treated with vehicle, mito-apocynin (mAPO, 1 μM, HY-135869, MedChemExpress), or cyclosporin A (CsA, 1 μM, C1832, Sigma) for 1 hour, and then with dexamethasone (DEX, 1 µM, D2915, Sigma) for 48 hours. For all conditions, primary neuronal cultures were collected for immunoblotting, mitochondrial function assay or immunostaining at 13-14 DIV.

### Media preparation for immunoblot and ELISA

When indicated, the cell culture media were prepared as described previously^26,27^. Briefly, the media was collected and concentrated using Pierce™ Protein Concentrators PES with 30K molecular-weight cutoff (Thermo Scientific, 88531) followed by centrifugation at 2000g for 20min. The supernatant was then subjected to sequential centrifugation steps: 30 min at 10,000 g, 30 min at 21,000 g, and finally 70 min at 100,000 g to deplete extracellular vesicles (EVs). The remaining supernatant was used for the experiments as indicated.

### ELISA

EV-depleted media samples (50 µL volume) were used for measurement of Tau concentration by mouse-specific total Tau ELISA kit (KMB7011, Thermo Scientific) according to manufacturer’s instructions.

### Immunoblotting

Protein extracts were separated by SDS/PAGE (10% Tris-Glycine gel; XP00105BOX, Invitrogen), then transferred to nitrocellulose membranes (10600001, Amersham). After blocking in TBST buffer (20 mM Tris-HCl, 150 mM sodium chloride, 0.1% Tween-20) containing 5% (wt/vol) nonfat dry milk for 1 h at room temperature, the membrane was incubated with primary antibodies overnight at 4°C, then with secondary antibodies for 1 h at room temperature. The following antibodies were used: AT8 (MN1020, ThermoFisher Scientific), PHF-1 (from Dr. Peter Davies), Tau5 (ab80579, Abcam), p-GR (4161S, Cell Signaling), GR (12041S, Cell Signaling), HSP70 (333800,Thermo Fisher), HSP90 (MA545102,Thermo Fisher), FKBP51 (PA1020, Thermo Fisher), anti-CypD (ab110324, Abcam), anti-KDM1/LSD1 (ab17721, Abcam), anti-Tom20 (OA241-4F3, NOVUS), anti-Tubulin (ab4074, Abcam). IRDye 800CW goat anti-mouse IgG secondary antibody (P/N: 926-32210, LI-COR), IRDye 680CW goat anti-rabbit IgG secondary antibody (P/N: 926-68071, LI-COR). Membranes were visualized by Odyssey Infrared Imager (model 9120, LI-COR Biosciences), and relative optical densities of bands determined by Fiji/ImageJ software. Full immunoblots used in the figures of this manuscript are shown in **Fig. S3**.

### Immunofluorescence staining of brain slices and cultured neurons

Floating brain sections or fixed primary neurons were incubated overnight with the following primary antibodies: mouse anti-oligomeric Tau antibody TOMA-1 (1:2500, Millipore sigma, MABN819), mouse anti-Synapsin I antibody (1:1000, 611393, BD Biosciences), mouse anti-phospho-Tau pSer202/Thr205 (1:1000, MN1020, ThermoFisher Scientific), and chicken MAP2 (1:5000, ab5392, Abcam). They were then incubated for 1 h with secondary antibodies (Alexa Fluor 488, 594, and 633 goat anti-rabbit or anti-mouse IgG, 1:2000 dilution). Coverslips were mounted with VectaShield (Vector Laboratories) and sealed with clear nail polish. Images were acquired with a 40X objective (Neofluar, NA 1.4) on a Zeiss LSM 800 confocal microscope running Zen2 software or with a 2X objective on an Eclipse 90i dual laser-scanning confocal microscope (NIKON, for lower magnification images in **Figures 1F, 8B**). The images were manually measured and quantified using the auto-threshold settings in Fiji/ImageJ software.

**Figure 1.**
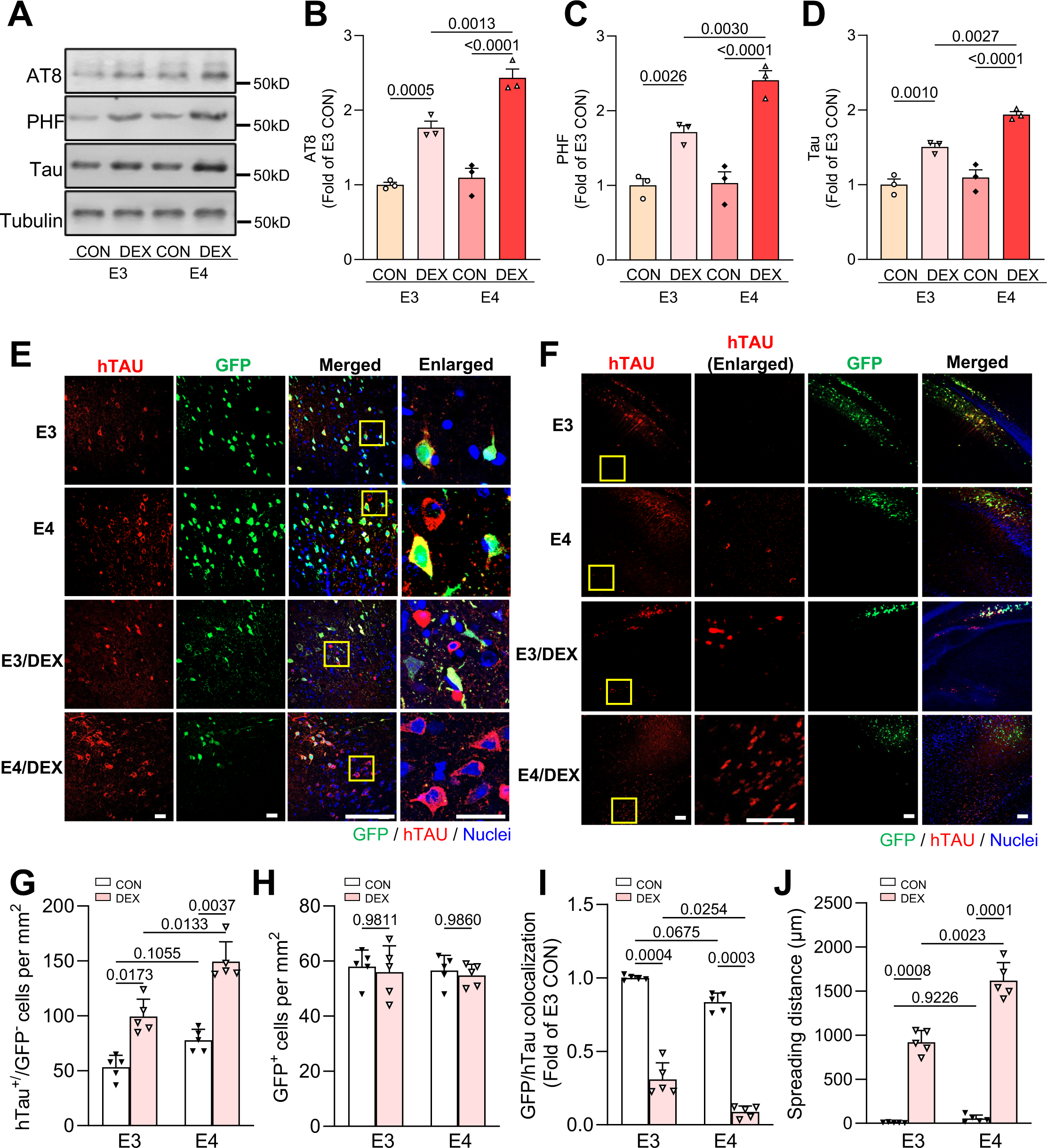
Glucocorticoid-induced tau phosphorylation and spreading are enhanced in E4 mice. (**A**–**D**) Representative immunoblots (**A**) and quantification (**B**–**D**) of AT8, PHF1, and total tau (Tau5) immunoreactivity in lysates from hippocampal tissue of 9-10-month-old E3/E4 mice treated with vehicle (CON) or dexamethasone (DEX). Intensity values are expressed relative to tubulin and normalized to the E3 CON condition (*P-*values indicated on graphs; data presented as mean ± SD; one-way ANOVA with Tukey’s multiple comparisons test; n=3 mice/condition for **B-D**). **(E)** Representative images showing hTau (red) and GFP (green) in CA1 neurons of mice treated as indicated. Nuclei are stained with DAPI (blue). The right column shows enlarged regions (indicated by yellow boxes). Scale bars, 50 µm. (**F**) Representative images depicting the spreading of hTau (red) from GFP^+^ cells near the injection site in mice treated as indicated. Yellow boxes indicate enlarged regions. Scale bars, 200 µm. (**G**) Quantification of hTau^+^/GFP^−^ cells per mm^2^ in mice treated as indicated. (**H**) Quantification of GFP^+^ cells per mm^2^ in mice treated as indicated. (**I**) Quantification of the GFP/hTau colocalization ratio in each condition, normalized to E3 CON condition. (**J**) Quantification of Tau spreading distance (μm) for each condition. For **G-J**, *P-*values are indicated on graphs; data presented as mean ± SD; two-way ANOVA with multiple comparisons test; n=5 mice/group. Each point represents an individual mouse.

### Mitochondrial/Cytosolic/Nuclear Isolation

Mitochondrial isolation was performed using the Mitochondrial extraction kit (Novus, NBP2-29448). Briefly, tissues were washed using ice-cold 1X PBS, then 5 ml of ice-cold homogenization buffer (provided by the kit) was added per gram of tissue. Following homogenization in a Dounce-type homogenizer, cell suspensions were transferred to tubes and nuclear pellets collected following centrifugation at 700g for 10 min at 4℃. The cytosolic fraction (supernatant) and the mitochondrial pellet were collected separately according to the manufacturer’s instructions.

### Mitochondria functional assays

Complex I activity and ATP production were measured from 13-14 DIV hippocampal neurons or hippocampal tissue with Complex I Enzyme Activity Microplate Assay Kit (ab109721, Abcam) and ATP Assay Kit (Colorimetric/Fluorometric, ab83355, Abcam), respectively, according to the manufacturer’s instructions.

### Evaluation of mitochondrial and cerebral reactive oxygen species

To estimate the production of reactive oxygen species (ROS), brain sections from mouse hippocampal tissue or primary neurons were exposed to 1 μM MitoSOX Red (M36008, ThermoFisher), a fluorochrome specific for anion superoxide produced in the inner mitochondrial compartment, at 37°C for 30 min. Images were acquired at 37°C with a 40X objective (Neofluar, NA 1.4) on a Zeiss LSM 800 confocal microscope running Zen2 software. Quantification of staining intensity and the percentage of area occupied by Mitosox was measured and quantified by using the auto-threshold settings in Fiji/ImageJ software.

Intracellular ROS levels were measured by election paramagnetic resonance (EPR) spectroscopy. Brain tissues were incubated with CMH (cyclic hydroxylamine 1-hydroxy-3-methoxycarbonyl-2, 2, 5, 5-tetramethyl-pyrrolidine, 100 μM) for 30 minutes, then washed three times with ice-cold PBS. The tissues were collected and homogenized with 100 μl of PBS for EPR measurement. EPR spectra were collected, stored, and analyzed via electron paramagnetic resonance (EPR, Bruker ESR 5000, Germany)^28^.

### Evaluation of mPTP opening

Hippocampal neurons (1 × 10^2^ cells/well, DIV 13-14) plated onto Lab-Tek 4-well chamber slides were treated with 1 μM Calcein Green AM (C3099, Thermo Fisher Scientific) at 37°C for 30 min, then treated ±1 mM cobalt chloride (CoCl^2^) for 30 min. Images were acquired at 37°C with a 40X oil-immersion objective (Neofluar, NA 1.3) on an epifluorescence microscope (Axio Observer Z1, Zeiss) with Colibri LED light source, EMCCD camera (Hamamatsu) and Zen 2012 (blue edition) software. Quantification of calcein fluorescence intensity was measured and quantified using the auto-threshold settings in Fiji/ImageJ software.

### AAV injection procedure

This procedure was described in detail in previous studies^26,27^. AAV.CBA.eGFP.2A.P301L-Tau plasmid (Addgene plasmid #140425) was packaged into AAV8 serotype. Prior to AAV injection, male/female mice (5/group) were administered DEX (5 mg/kg, i.p.injection) or mito-apocynin (3 mg/kg, i.p. injection) for 7 days. Stereotactic AAV injections were performed under standard aseptic surgery conditions as previously described^27,29^. Briefly, mice were anaesthetized with isoflurane (2%), placed in a stereotactic frame (digital stereotaxic device, Stoelting Co.), and injected bilaterally with 2 μl of AAV in hippocampal region CA1 (at the following coordinates relative to Bregma: A/P −2.7 mm, M/L ±2 mm, D/V −1.5 mm) with a 10 μl Hamilton syringe at a rate of 0.25 μl/min by a Nano-injector system (Stoelting microsyringe pump, Stoelting Co.). Afterwards, the skin over the injection site was sutured and mice were placed on a warming pad during their recovery. Mice were then administered DEX or mito-apocynin for an additional 14 days prior to euthanasia. Control animals received daily i.p. injections of PBS (DEX vehicle) or 15% PEG400 in PBS (mito-apocynin vehicle).

### Quantification of Tau spreading

hTau^+^ neurons (detected by immunostaining with Tau13 antibody) were counted in hippocampi of coronal brain sections near the site of AAV injection, identified by the dense cluster of GFP^+^ neurons. Tau spreading was quantified as in our previous studies^27,29,30^, by calculating the number of hTau^+^ neurons in the hippocampus that did not exhibit GFP fluorescence (hTau^+^/GFP^-^ neurons) per mm^2^ and the fraction of hTau^+^ cells that were GFP^+^ (GFP/hTau colocalization). For each condition, we also measured the maximum distance between hTau^+^ neurons in the vicinity of the hippocampal formation and the cluster of GFP^+^ neurons near the injection site, using the Fiji/Image J measurement tools.

### Statistical analysis

All values were expressed as the mean ± SEM. All graphing and statistical analyses were performed using GraphPad Prism (GraphPad Prism10.Ink). Statistical details of experiments are provided in the figure legends. Statistical analyses were performed with unpaired, two-tailed t-test or one-way ANOVA with Tukey’s test for multiple comparisons. Values of *p* < 0.05 were considered statistically significant. Investigators were blinded to treatment conditions when performing analyses for all experiments.

## Data Availability

The data generated and analyzed in this study are available from the corresponding author upon reasonable request.

## Results

### Glucocorticoid-induced tau pathology is amplified by *APOE4*

To determine whether the *APOE4* genetic variant exacerbates GC-induced brain pathology, we first assessed tau accumulation and phosphorylation in middle-aged (8-9-month-old) *APOE3* (E3) and *APOE4* (E4) humanized knockin mice. Animals were administered vehicle or the synthetic GC dexamethasone (DEX; 5 mg/kg daily, i.p. injection) for 15 days, a widely-used treatment that mimics the high circulating GC levels induced by chronic stress^25,31-34^. DEX administration had similar effects on body weight loss and suppression of endogenous corticosterone levels in both genotypes (Fig. **S1A-B**), indicative of their equivalent physiological responses to GCs. Following brain tissue harvest, we measured the levels of total tau and phospho-tau epitopes commonly associated with tauopathies such as Alzheimer’s by immunoblotting hippocampal lysates with Tau5 (total tau), PHF-1 (S396/S404), and AT8 (S202/T205) antibodies. We saw no difference in total or phosphorylated tau levels in lysates from vehicle-treated E4 versus E3 mice (Fig. **1A-D**), consistent with a previous report that E4 targeted replacement animals lack overt tau pathology at 9 months of age^35^. DEX treatment significantly increased total and phospho-tau levels in both E3 and E4 genotypes, but caused a more dramatic increase in tau accumulation in the E4 hippocampi (∼2.5-fold vs. ∼1.5-fold; Fig. **1A-D**). Since tau phosphorylation is known to stimulate its secretion^36,37^, and we recently found that GCs promote tau secretion and spreading^38^, we also evaluated GC-induced tau spreading in E3 and E4 mice. Here, 9-10-month-old animals were pre-treated with vehicle or DEX for 7 days, then injected in hippocampal area CA1 with adeno-associated virus (AAV) co-expressing GFP and the human frontotemporal dementia-associated tau mutant P301L (hTau) separated by the 2A cleavage sequence (AAV.CBA.eGFP.2A.P301L-Tau)^30^, enabling visualization of trans-cellular tau spreading. Animals were then administered vehicle or DEX for 14 more days, followed by brain tissue harvest and immunostaining with antibodies against hTau (Fig. **1E-F**). Tau propagation was quantified as previously described^29,30^, by calculating hTau^+^/GFP^-^ neurons per mm^2^, GFP/hTau colocalization, and maximal spreading distance of hTau from GFP^+^ neurons (Fig. **1G-J**). These studies revealed that while AAV transduction efficiency (number of GFP^+^ cells per mm^2^) was equivalent between genotypes (Fig. **1H**), tau propagation was significantly enhanced in DEX-treated E4 versus E3 animals, based on the increased number of hTau^+^/GFP^-^ neurons, decreased GFP/hTau colocalization, and increased hTau spreading distance in E4 hippocampus (∼1500 μm vs. ≤1000 μm in the E3 background; Fig. **1G, I****-J**). Together, these findings demonstrate that GC-induced tau accumulation and spreading is significantly exacerbated by the *APOE4* variant.

### Underlying mitochondrial dysfunction in E4 animals augments the impact of GCs

We recently showed that GC-induced tau pathology is precipitated by mitochondrial damage^17^. ApoE4 expression is associated with mitochondrial dysfunction in humans, mice, and cell culture models^18-21,35,39-41^, and may enhance mitochondrial susceptibility to GC-mediated damage, thus promoting more extensive downstream tau pathology. To investigate this possibility, we first measured reactive oxygen species (ROS) levels, an indicator of mitochondrial dysfunction, using highly-sensitive electron paramagnetic resonance (EPR) spectroscopy on hippocampal tissue of the 9-10-month-old vehicle- and DEX-treated E3 and E4 animals described above (Fig. **1**). We saw a nearly two-fold increase in ROS levels in control E4 versus E3 tissue (Fig. **2A-B**), indicative of underlying oxidative stress in E4 hippocampi. Moreover, DEX treatment increased ROS to a greater extent in the E4 samples, suggesting mitochondrial vulnerability to GCs. To follow up on these findings, we evaluated mitochondrial function in this tissue more directly by measuring the activity of complex I, the first component of the mitochondrial electron transport chain, and ATP production. Interestingly, both complex I activity and ATP production were significantly reduced in E4 compared to E3 tissue under control conditions, and both were inhibited to a greater extent by DEX treatment in the E4 background (Fig. **2C-D**). We further monitored the impact of DEX on mitochondrial ROS production in E3 and E4 hippocampi using the fluorescent superoxide sensor MitoSOX. Here, we found that mROS levels were higher at baseline in E4 vs. E3 hippocampal tissue, and also more dramatically increased by DEX (Fig. **2E-F**). Since we previously showed that oligomeric tau is closely associated with mROS in hippocampal neurons^17^, we also measured oligomeric tau levels in these hippocampal slices with TOMA-1 antibodies (Fig. **2E, G**). Similar to its impact on mROS, we found that DEX treatment increased oligomeric tau to a greater degree in the E4 versus the E3 background, and that this oligomeric tau colocalized with mROS in neuronal cell bodies (Fig. **2E, G**). These data suggest that underlying mitochondrial dysfunction aggravates GC-driven mitochondrial damage and subsequent tau oligomerization in E4 animals.

**Figure 2.**
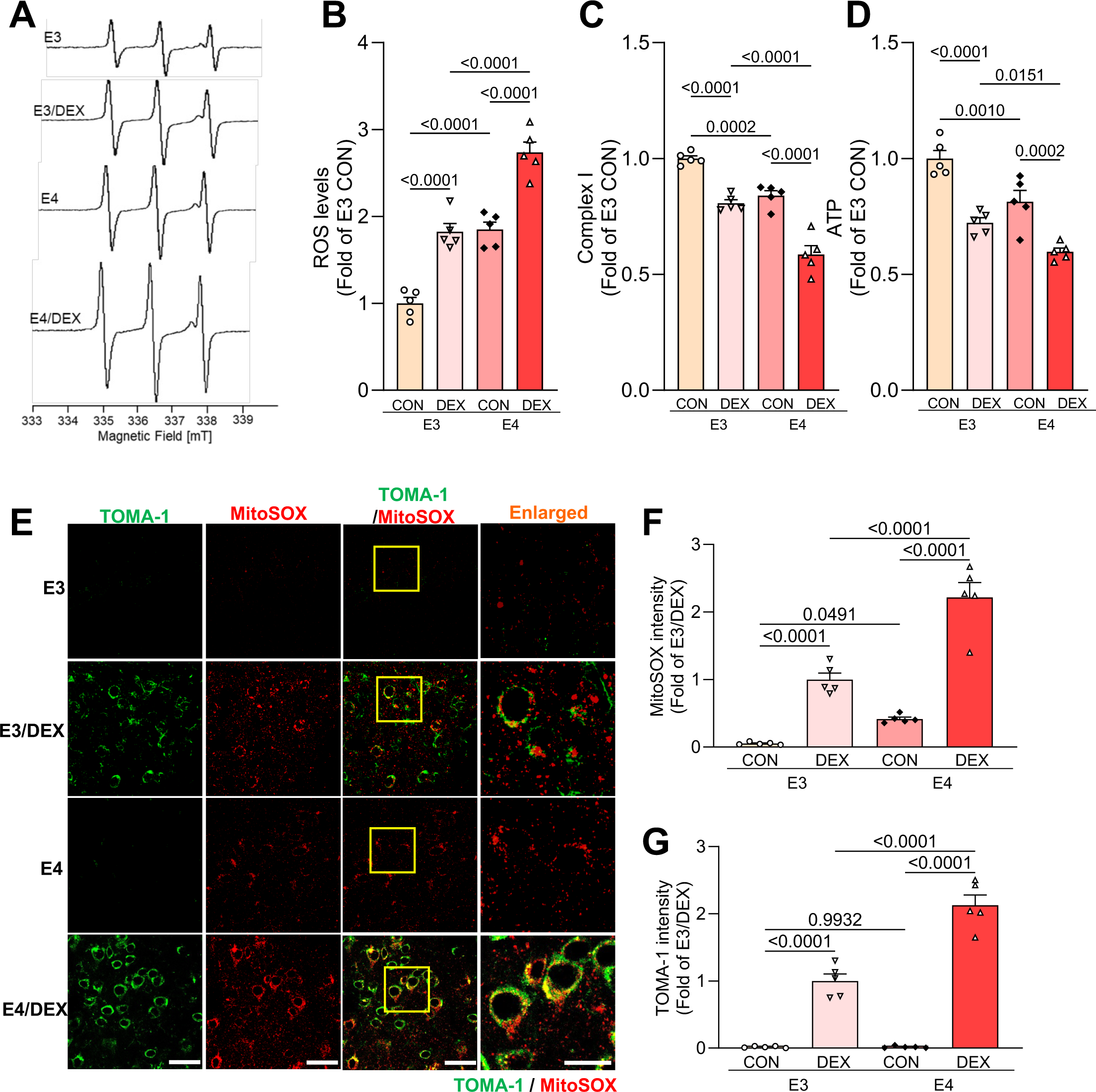
Underlying mitochondrial dysfunction amplifies the effect of GCs in E4 mice. (**A-B**) Representative spectra from electron paramagnetic resonance (EPR) spectroscopy of hippocampal tissue from 9-10-month-old E3 and E4 mice treated with vehicle (CON) or dexamethasone (DEX)(**A**), and accompanying quantification (**B**). Peaks in the spectra represent ROS levels (*P-*values indicated on graphs; data presented as mean ± SD; unpaired *t*-test; n=5 mice/condition). (**C-D**) Complex I activity (**C**) and ATP levels (**D**) in hippocampal tissue from E3/E4 mice treated as indicated, normalized to the E3 CON condition (*P-*values indicated on graphs; data presented as mean ± SD; one-way ANOVA with Tukey’s multiple comparisons test; n=5 mice/condition). (**E**–**G**) Representative images (**E**) and quantification (**F**, **G**) of MitoSOX and TOMA-1 fluorescence intensity in slices from hippocampal area CA1 of mice treated as indicated. Right-most column shows enlarged regions (indicated by yellow boxes). Scale bars=50 µm. Intensity values are normalized to the E3 DEX condition (*P-*values indicated on graphs; data presented as mean ± SD; one-way ANOVA with Tukey’s multiple comparisons test; n=5 mice/condition). Each point represents an individual mouse.

### GR activation is enhanced in *APOE4* carriers

GCs signal through their binding to glucocorticoid receptors (GRs), ligand-dependent transcription factors that regulate gene expression via binding to glucocorticoid response elements of target genes^42^. GRs are primarily localized to the cytoplasm of cells, but in response to GC binding they translocate to the nucleus and/or mitochondria to mediate transcription of nuclear and/or mitochondrial DNA^42^. GR ligand binding and activation are regulated by interactions with its molecular chaperones Hsp90, Hsp70, and FKBP51/52^42,43^. In particular, Hsp90 has a role in GR activation^43,44^ and Hsp90 mRNA was found to be upregulated in neurons and glia carrying *APOE4^23^*, suggesting that ApoE4 expression may enhance GR activation and signaling. To explore this concept, we measured the levels of plasma corticosterone, the endogenous rodent GC, by ELISA, and of Hsp90, Hsp70, and FKBP51 by immunoblotting hippocampal lysates from 9-10-month-old E3 and E4 mice. Indeed, while corticosterone levels were indistinguishable between genotypes, levels of all three co-chaperones were significantly elevated in E4 versus E3 tissue (Fig. **3A-C**), suggesting that GR activation could be altered in E4 brains. As a proxy for GR activation/signaling, we next examined GR levels in cytoplasmic, nuclear, and mitochondrial fractions isolated from hippocampal tissue of these animals. Consistent with the other findings, we observed significantly increased nuclear and mitochondrial GR levels, and concomitantly decreased cytoplasmic GR levels, in lysates from E4 compared to E3 animals (Fig. **3D-E**), indicative of increased GR activation and translocation into the nucleus and mitochondria. We further measured GR phosphorylation, another indicator of GR activation, in hippocampal lysates from the DEX-treated E3 and E4 mice. We found that phospho-GR levels were more significantly elevated by DEX in E4 hippocampus (Fig. **3F-G**), consistent with the concept that ApoE4 induces cellular changes that lower the threshold of activation for GR.

**Figure 3.**
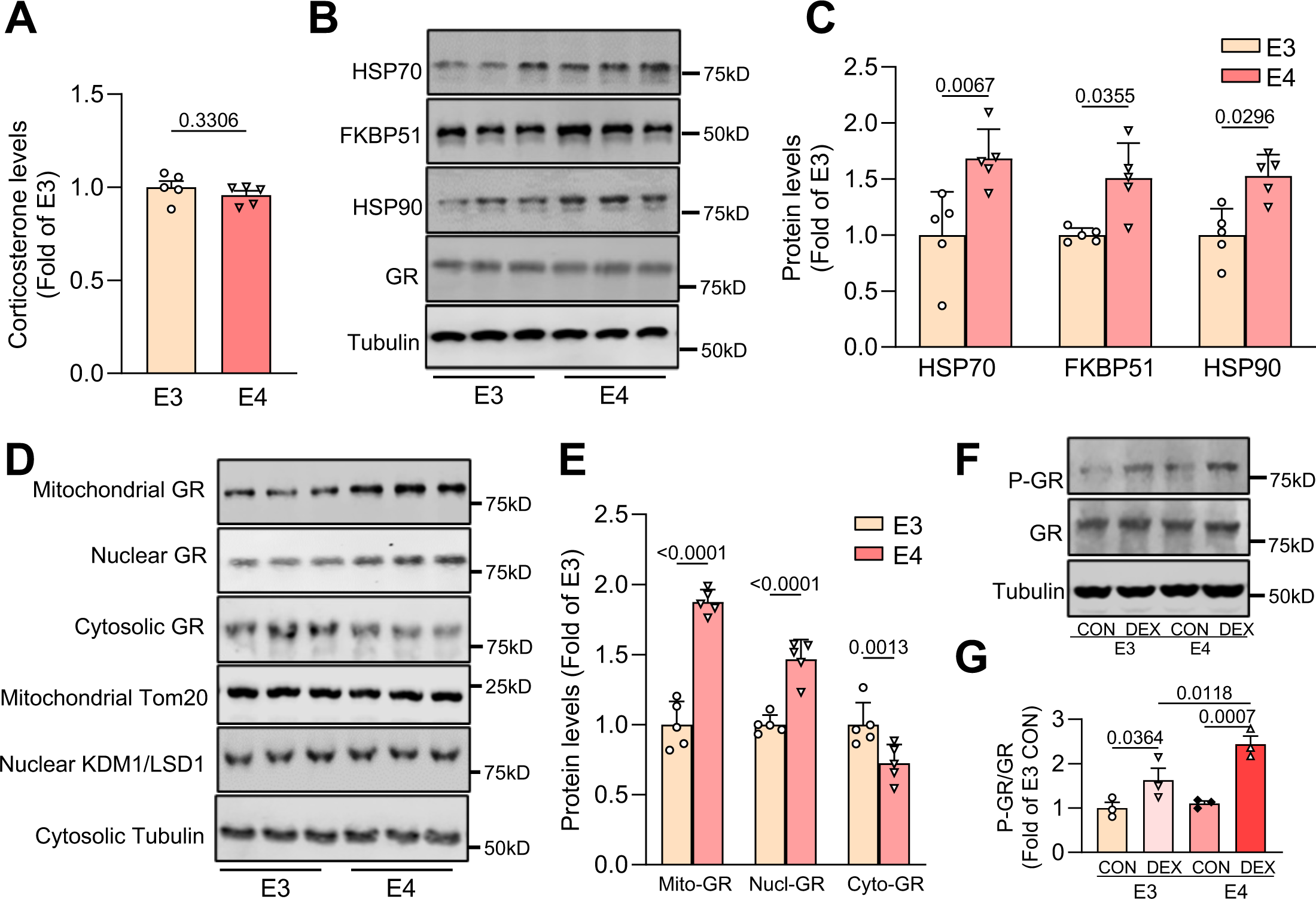
GR activation is enhanced in hippocampal tissue from E4 animals. (**A**) Quantification of corticosterone levels in plasma from 9-10-month-old E3/E4 mice, measured by ELISA (*P-*values indicated on graphs; data presented as mean ± SD; unpaired *t*-test; n=5 mice/condition). (**B-C**) Representative immunoblots (**B**) and quantification (**C**) of Hsp70, FKBP51, and Hsp90 immunoreactivity in hippocampal lysates from the indicated mice. Intensity values are expressed relative to tubulin and normalized to the E3 group (*P-*values indicated on graphs; data presented as mean ± SD; unpaired *t*-test; n=5 mice/condition). (**D**–**E**) Representative immunoblots (**D**) and quantification (**E**) of mitochondrial, nuclear, and cytosolic GR, based on immunoreactivity from mitochondrial, cytosolic, and nuclear fractions of the indicated mice. Intensity values are expressed relative to mitochondrial marker TOM20, nuclear marker KDM1/LSD1, and cytosolic marker tubulin, and normalized to the E3 group (*P-*values indicated on graphs; data presented as mean ± SD; unpaired *t*-test; n=5 mice/condition). (**F**–**G)** Representative immunoblots (**F**) and quantification (**G**) of phospho-GR levels, normalized to total GR, in hippocampal lysates from 9-10-month-old E3/E4 mice +/- dexamethasone (DEX). Intensity values are expressed relative to tubulin and normalized to the E3 CON group (*P-*values indicated on graphs; data presented as mean ± SD; one-way ANOVA with Tukey’s multiple comparisons test; n=3 mice/condition). Each point represents an individual mouse.

To further investigate whether ApoE4 augments neuronal responsiveness to GC/GR signaling, we treated 14 day *in vitro* (DIV) primary hippocampal neurons from E3 and E4 mice for 48 hours with 200 nM DEX, a concentration five times lower than that which we typically use for our *in vitro* experiments (1 μM). Treatment with 200 nM DEX did not alter phospho-GR levels in E3 neurons, but increased these levels by three-fold in E4 neurons (Fig. **4A-B**). Similarly, 200 nM DEX impaired complex I activity and ATP production in E4 but not E3 neurons (Fig. **S2A-B**), and induced the accumulation of phospho- and total tau in E4 neurons only, as measured by immunoblotting of neuronal lysates with AT8, PHF1, and Tau5 antibodies (Fig. **4C-F**). This low DEX concentration also promoted tau secretion from E4 neurons only, based on immunoblotting and ELISA of concentrated media harvested from E3 and E4 neurons (Fig. **4G-K**). The increase in tau secretion was not due to compromised health or plasma membrane integrity of E4 neurons, as the levels of lactate dehydrogenase (LDH), a cytosolic protein, were similar in media from E4 vs. E3 neurons and not altered by DEX treatment (Fig. **4K**). These findings indicate that GC/GR signaling and downstream tau pathogenesis are enhanced in the presence of ApoE4.

**Figure 4.**
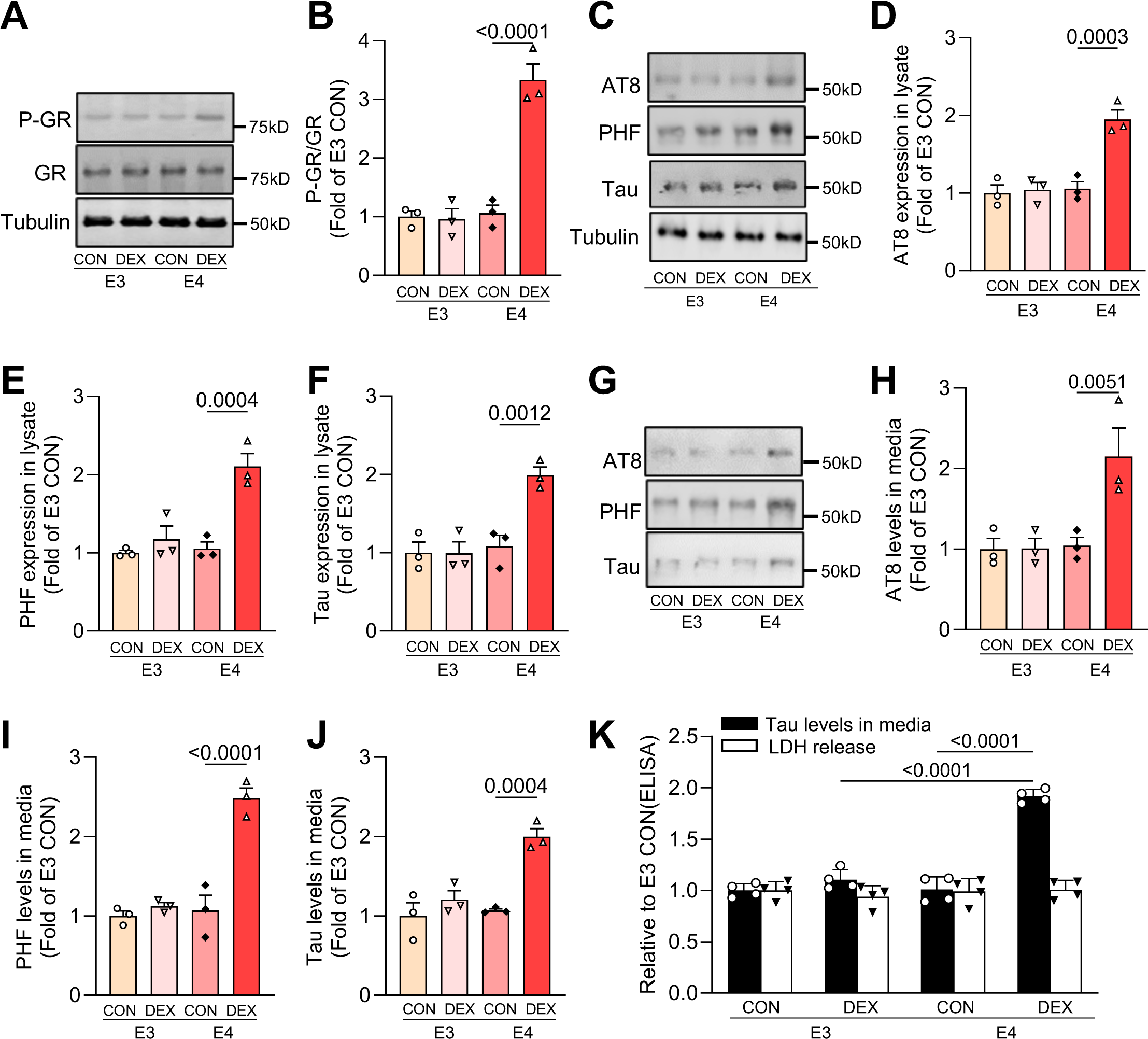
Low GC levels induce tau pathogenesis in E4 but not E3 hippocampal neurons. **(A–B**) Representative immunoblots (**A**) and quantification (**B**) of phospho-GR levels, normalized to total GR, in lysates from 14 DIV E3/E4 hippocampal neurons treated with vehicle (CON) or dexamethasone (DEX). Intensity values are expressed relative to GR and normalized to the E3 CON group (*P-*values indicated on graphs; data presented as mean ± SD; one-way ANOVA with Tukey’s multiple comparisons test; n=3 samples/condition). (**C**–**F**) Representative immunoblots (**C**) and quantification (**D**–**F**) of AT8, PHF1, and total tau (Tau5) immunoreactivity in lysates from the indicated treatment conditions. Intensity values are expressed relative to tubulin and normalized to the E3 CON condition (*P-*values indicated on graphs; data presented as mean ± SD; one-way ANOVA with Tukey’s multiple comparisons test; n=3 samples/condition). (**G**–**J)** Representative immunoblots (**G)** and quantification (**H**–**J)** of AT8, PHF1, and total tau (Tau5) immunoreactivity in extracellular vesicle (EV)-depleted media from the indicated treatment conditions. Intensity values are normalized to the E3 CON condition (*P-*values indicated on graphs; data presented as mean ± SD; one-way ANOVA with Tukey’s multiple comparisons test; n=3 samples/condition). (**K)** Quantification of ELISA for total tau levels (black bars) and LDH release (white bars) in extracellular vesicle (EV)-depleted media from the indicated treatment conditions (*P-*values indicated on graphs; data presented as mean ± SD; one-way ANOVA with Tukey’s multiple comparisons test; n=4 samples/condition).

### mPTP opening drives GC-induced tau pathogenesis in E4 neurons

We previously showed that GCs induce mitochondrial damage by promoting the opening of the mitochondrial permeability transition pore (mPTP), a channel on the inner mitochondrial membrane comprising several components including the F^1^/F^0^ ATP synthase and the mitochondrial matrix protein cyclophilin D (CypD)^45^. Activation/opening of the mPTP is triggered by CypD^45^, and we showed that GCs promote this event by transcriptionally upregulating CypD, leading to mitochondrial depolarization, overproduction of reactive oxygen species (ROS), and tau phosphorylation and oligomerization in hippocampal neurons^17^. To determine whether this mechanism is responsible for mitochondrial damage in ApoE4 neurons, we measured mPTP opening and CypD levels in 14 DIV E3 and E4 hippocampal neurons treated with vehicle or 200 nM DEX for 48 hours. mPTP opening was assessed via live imaging with the Co^2+^-calcein assay^46^, in which CoCl^2^-mediated quenching of calcein dye occurs upon pore opening, and CypD levels were assessed via immunoblotting of hippocampal lysates. In E3 neurons, 200 nM DEX treatment did not elicit CoCl^2^-mediated calcein quenching (Fig. **5A-B**) nor the upregulation of CypD (Fig. **5C-D**), demonstrating that this DEX concentration does not trigger mPTP opening. In contrast, E4 neurons treated with DEX exhibited significant CoCl^2^-mediated quenching (∼50%; Fig. **5A-B**) and a two-fold increase in CypD expression (Fig. **5C-D**), showing that even this low DEX concentration upregulates CypD and induces mPTP opening in the *APOE4* background. Since we previously found that the CypD inhibitor cyclosporin A (CsA) and the mitochondrially-targeted NADPH oxidase inhibitor mito-apocynin (mAPO) prevented GC-induced mPTP opening and downstream mitochondrial dysfunction and tau pathology in wild-type neurons^17^, we tested whether these compounds had the same impact in E4 neurons. Indeed, while 200 nM DEX caused a 50% reduction in complex I activity and ATP production, both CsA (1 μM) and mAPO (1 μM) prevented these effects (Fig. **5E-F**) as well as the DEX-induced accumulation of total and phospho-tau in E4 neurons (Fig**. 5G-J**). Further, these drugs inhibited DEX-induced tau oligomerization and mROS production (Fig. **5K-L**; Fig. **S2C-D**), demonstrating their protective role against GC-driven mitochondrial dysfunction and tau pathogenesis in E4 neurons.

**Figure 5.**
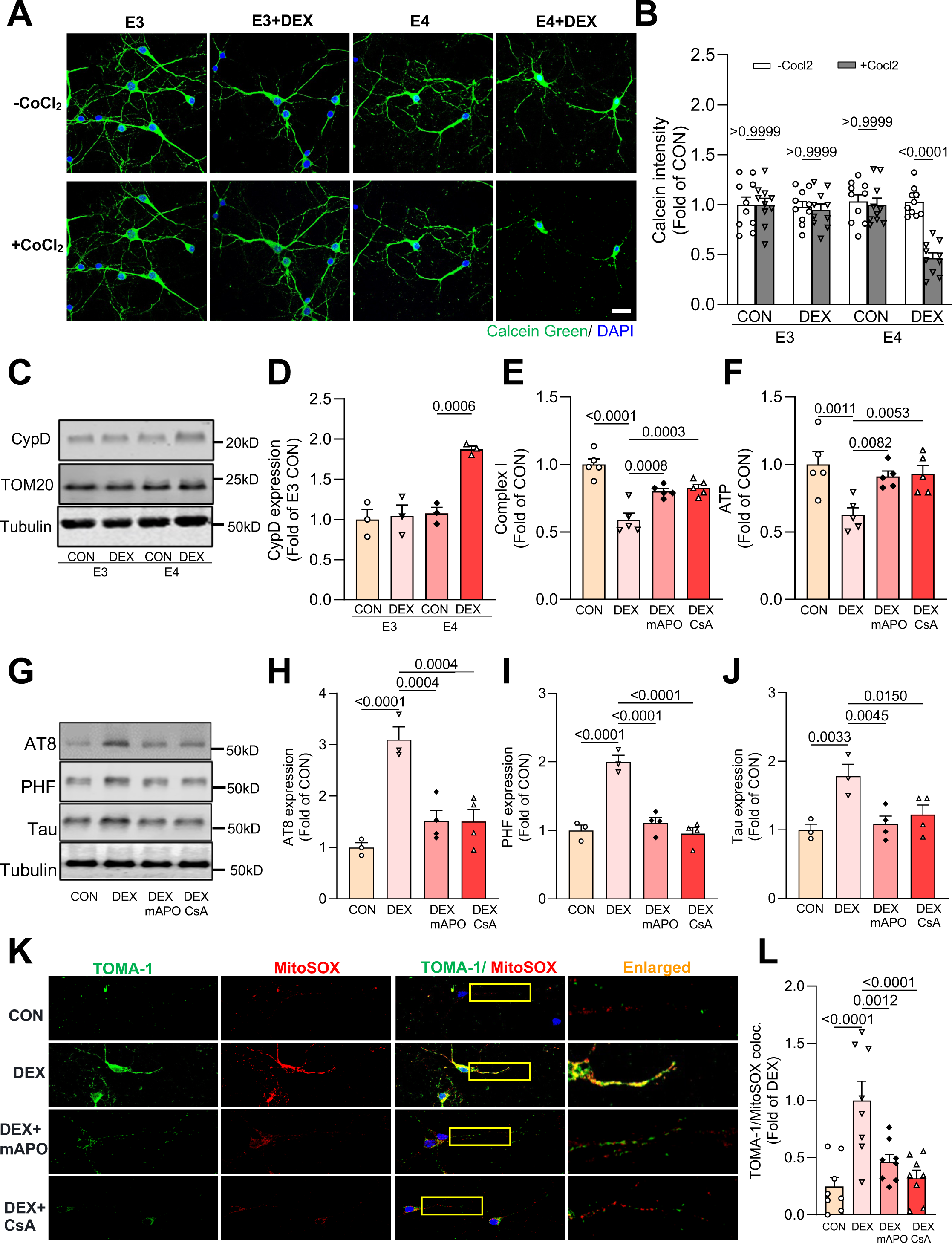
mPTP opening drives GC-induced mitochondrial dysfunction and tau pathology in E4 neurons. (**A**-**B**) Representative images (**A**) and quantification (**B**) of calcein green fluorescence intensity with or without CoCl^2^ in E3/E4 hippocampal neurons treated with vehicle (CON) or dexamethasone (DEX). Scale bar = 25 µm. Intensity values are normalized to the CON condition (*P*-values indicated on graph; data presented as mean ± SD; two-way ANOVA with multiple comparisons test; n=10 fields of view/condition). (**C-D**) Representative immunoblots (**C**) and quantification (**D**) of CypD and TOM20 immunoreactivity in lysates from E3/E4 hippocampal neurons treated as indicated. Intensity values are expressed relative to tubulin and normalized to the E3 CON condition (*P-*values indicated on graphs; data presented as mean ± SD; one-way ANOVA with Tukey’s multiple comparisons test; n=3 samples/condition). (**E-F**) Complex I activity (**E**) and ATP levels (**F**) in E4 hippocampal neurons treated with vehicle (CON), dexamethasone (DEX), DEX + mito-apocynin (mAPO) and DEX + cyclosporin A (CsA), normalized to the CON condition (*P-*values indicated on graphs; data presented as mean ± SD; one-way ANOVA with Tukey’s multiple comparisons test; n=5 samples/condition). (**G-J**) Representative immunoblots (**G**) and quantification (**H**-**J**) of AT8, PHF1, and total tau (Tau5) immunoreactivity in cell lysates from E4 hippocampal neurons treated as indicated. Intensity values are expressed relative to tubulin and normalized to the CON condition (*P-*values indicated on graphs; data presented as mean ± SD; one-way ANOVA with Tukey’s multiple comparisons test; n=3-4 samples/condition). (**K**–**L**) Representative images (**K**) and quantification of TOMA-1 co-localization with MitoSOX (**L**) in E4 hippocampal neurons treated as indicated. Enlarged regions (indicated by yellow boxes) are shown in the right column. Scale bars = 50 µm (*P-*values indicated on graphs; data presented as mean ± SD; one-way ANOVA with Tukey’s multiple comparisons test; n=8 fields of view/condition for **L**).

### Inhibition of mPTP opening is protective against tau pathogenesis in E4 animals

The *APOE4* variant is associated with greater risk for Alzheimer’s disease and other tauopathies compared to *APOE2* or *APOE3*^47,48^, and *APOE4* similarly exacerbates tau pathology and tau-related neurodegeneration in PS19 transgenic mice expressing the human frontotemporal dementia-linked P301S tau mutation^22^. To test whether mPTP opening plays a role in tau pathogenesis in *APOE4* carriers even in the absence of stress/high GC levels, we crossed E3 and E4 mice with PS19 tauopathy mice (hereafter called TE3 and TE4) and administered vehicle or mAPO (3 mg/kg; daily i.p. injection) to 1-1.5-month-old animals prior to the onset of any pathological changes. Ten weeks later, we euthanized the animals, harvested their brain tissue, and assessed tau-related pathology in the hippocampus. In hippocampal tissue, we observed a two-fold increase in phosphorylated tau levels in samples from TE4 versus TE3 animals (Fig. **6A-D**), as well as dramatically elevated oligomeric tau and associated mROS in neurons of area CA1 (Fig. **6E-G**). Administration of mAPO significantly attenuated these phenotypes (Fig. **6A-G**). Phosphorylated and oligomeric tau redistributes from axons into the somatodendritic compartment, inducing synaptic and neuronal loss that is thought to underlie cognitive decline in Alzheimer’s and other tauopathies^49,50^. To determine whether mAPO also mitigates this downstream impact of tau pathology, we examined synapse density in hippocampal area CA1. Immunostaining with antibodies against the presynaptic marker synaptophysin and the somatodendritic marker MAP2 revealed that the density of synapses and MAP2^+^ dendrites were significantly decreased in control TE4 versus TE3 animals, and both phenotypes were prevented by mAPO (Fig. **6H-J**). As anticipated, treatment with mAPO also normalized mitochondrial dysfunction in TE4 animals (Fig. **S2E-F**). These findings indicate that mitochondrial dysfunction, and mPTP opening in particular, plays a key role in ApoE4-mediated tau pathogenesis and associated synapse loss in the hippocampus.

**Figure 6.**
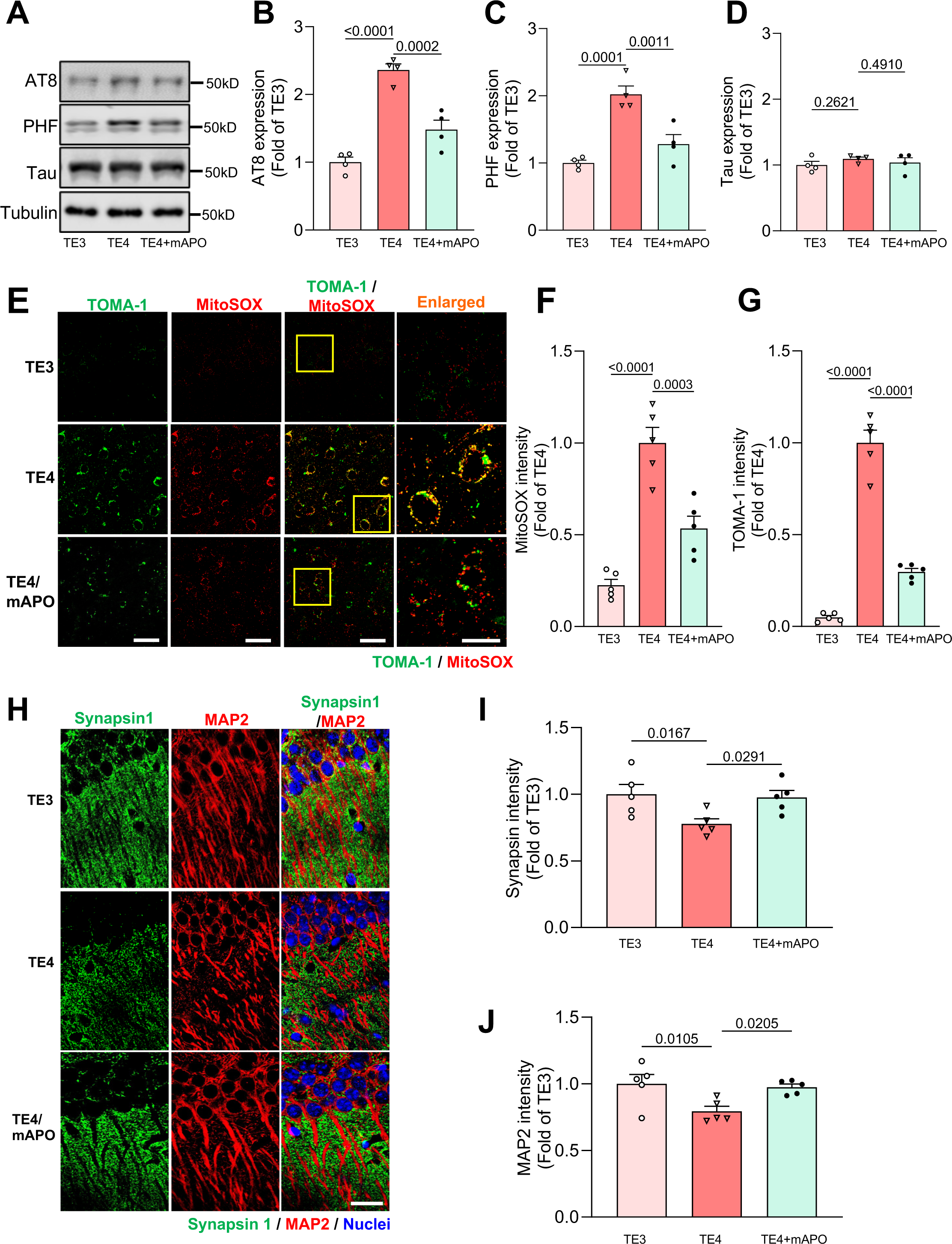
Mito-apocynin protects against mitochondrial dysfunction and tau pathology in TE4 animals. (**A**–**D**) Representative immunoblots (**A**) and quantification (**B**–**D**) of AT8, PHF1, and total tau (Tau5) immunoreactivity in hippocampal lysates from 3.5-4-month-old TE3 and TE4 mice after treatment with vehicle or mito-apocynin (mAPO). Intensity values are expressed relative to tubulin and normalized to the TE3 condition (*P-*values indicated on graphs; data presented as mean ± SD; one-way ANOVA with Tukey’s multiple comparisons test; n=4 samples/condition). (**E** -**F**) Representative images (**E**) and quantification of MitoSOX (**F**) and TOMA-1 (**G**) fluorescence intensity in hippocampal area CA1 of mice treated as indicated. Enlarged regions are shown in the right column (indicated by yellow boxes). Scale bars = 50 µm. Intensity values are normalized to TE4 controls given the absence of signal in the TE3 condition (*P-*values indicated on graphs; data presented as mean ± SD; one-way ANOVA with Tukey’s multiple comparisons test; n=5 mice/condition). (**H**–**J**) Representative images (**H**) and quantification (**I-J**) of Synapsin1a (green) and MAP2 (red) immunofluorescence intensity in hippocampal area CA1 of mice treated as indicated. DAPI (blue) labels nuclei. Scale bars = 50 μm. Synapsin1a and MAP2 intensity values are normalized to the TE3 condition (*P-*values indicated on graphs; data presented as mean ± SD; one-way ANOVA with Tukey’s multiple comparisons test; n=5 mice/condition). Each point represents an individual mouse.

While the previous experiment shows that mAPO confers remarkable protection against ApoE4-induced tau pathology, it was conducted in a mouse model overexpressing pathogenic P301S tau, a non-physiological condition. To study the efficacy of mAPO in a condition more relevant to human Alzheimer’s disease, we utilized aging E3 and E4 animals. In particular, we administered vehicle or mAPO (3 mg/kg, i.p. injection) to 15-16-month-old E3 and E4 animals for 15 days, then harvested brain tissue to measure tau phosphorylation/oligomerization and mitochondrial function as described above. In contrast to 9-10-month-old E4 (Fig. **1**) and 15-16-month-old E3 mice, the 15-16-month-old E4 animals exhibited a two-fold increase in the levels of phosphorylated tau in hippocampal tissue as measured by AT8 and PHF1 antibodies, with no change in total tau (Fig. **7A-D**). Treatment with mAPO had no impact on total or phospho-tau levels in E3 brain tissue, but normalized phospho-tau levels in E4 tissue to those of the E3 controls (Fig. **7A-D**). Similarly, complex I activity and ATP production in aging E4 animals were reduced by 50% compared to E3 animals, and mAPO normalized both of these to the level of E3 control animals without impacting baseline mitochondrial function in the E3 background (Fig. **7E-F**). Levels of mROS and oligomeric tau were likewise dramatically increased in aging E4 vs. E3 animals, and rescued in E4 animals by mAPO (Fig. **7G-I**).

**Figure 7.**
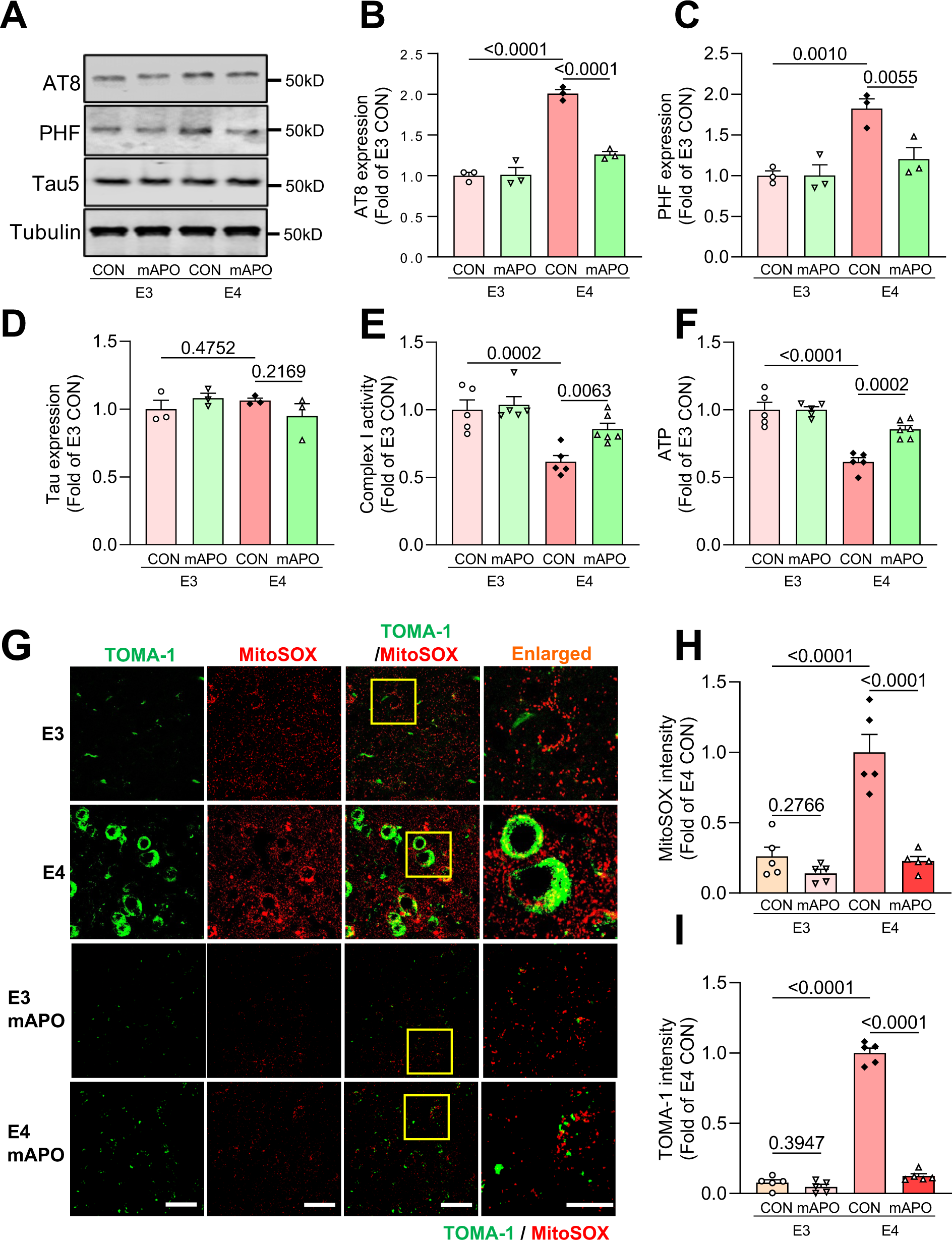
Mito-apocynin prevents mitochondrial dysfunction and tau pathology in aging E4 animals. (**A**–**D**) Representative immunoblots (**A**) and quantification (**B**–**D**) of AT8, PHF1, and total tau (Tau5) immunoreactivity in hippocampal lysates from 15-16-month-old E3 and E4 mice treated for 15 days with vehicle (CON) or mito-apocynin (mAPO). Intensity values are expressed relative to tubulin and normalized to the E3 CON condition (*P-*values indicated on graphs; data presented as mean ± SD; one-way ANOVA with Tukey’s multiple comparisons test; n=3 mice/condition). (**E -F**) Complex I activity (**E**) and ATP levels (**F**) in hippocampal tissues treated as indicated, normalized to the E3 CON condition (*P-*values indicated on graphs; data presented as mean ± SD; one-way ANOVA with Tukey’s multiple comparisons test; n=5-6 mice/condition). (**G**–**I**) Representative images (**G**) and quantification (**H** and **I**) of MitoSOX and TOMA-1 fluorescence intensity in hippocampal area CA1 of mice treated as indicated. Enlarged regions are shown in the right column (indicated by yellow boxes). Scale bars=50 µm. Intensity values are normalized to the E4 CON condition (*P-*values indicated on graphs; data presented as mean ± SD; one-way ANOVA with Tukey’s multiple comparisons test; n=5 mice/condition).

Finally, we examined the impact of mAPO on tau propagation, shown to be enhanced in the *APOE4* genotype^51,52^. Here, 15-16-month-old animals were administered vehicle or mAPO for one week prior to AAV injection of AAV.CBA.eGFP.2A.P301L-Tau in hippocampal area CA1, and tau spreading was assessed two weeks later as described above. In contrast to tau spreading in the younger (9-10-month-old) mice, for which no significant difference was observed between genotypes (Fig. **1**), spreading in 15-16-month-old mice was significantly increased for E4 versus E3 animals (Fig. **8A-B**). Specifically, E4 hippocampi had increased numbers of hTau^+^/GFP^-^ neurons per mm^2^, decreased GFP/hTau colocalization, and increased hTau spreading distance (>1000 μm for E4 vs. <100 μm for E3) compared to E3 hippocampi (Fig. **8C, E****-F**), while AAV transduction efficiency (GFP^+^ cells/mm^2^) was equivalent between genotypes (Fig. **8D**). Administration of mAPO treatment completely normalized the enhanced spreading in E4 brains, demonstrating that mitochondrial dysfunction, and mPTP opening in particular, plays an important role in driving tau propagation in *APOE4* carriers.

**Figure 8.**
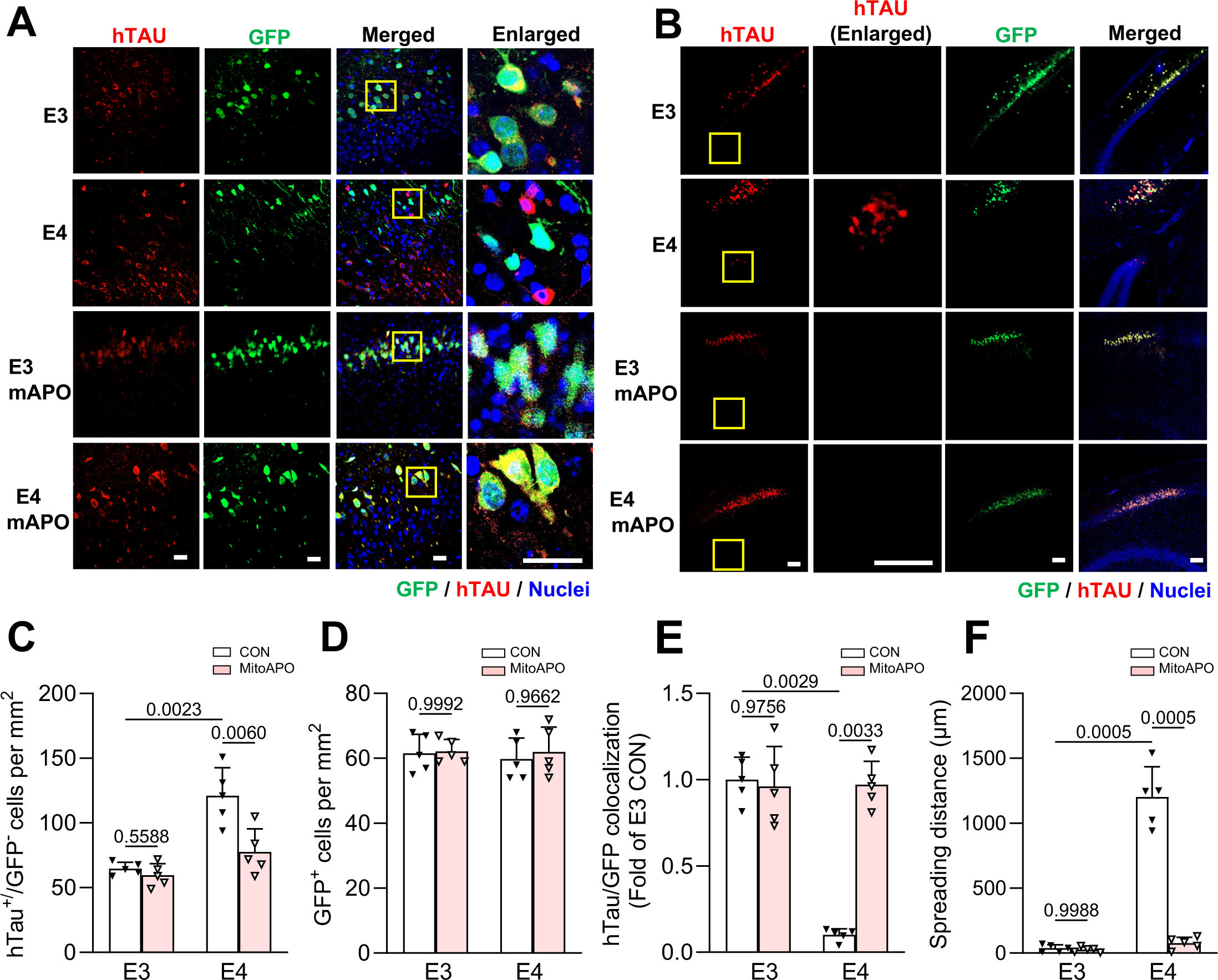
Mito-apocynin prevents tau spreading in aging E4 animals. (**A**) Representative images showing hTau (red) and GFP (green) in CA1 neurons of from 15-16-month-old E3 and E4 mice treated with vehicle (CON) or mito-apocynin (mAPO). Nuclei are stained with DAPI (blue). The right column shows enlarged regions (indicated by yellow boxes). Scale bars, 50 µm. (**B**) Representative images depicting the spreading of hTau (red) from GFP^+^ cells near the injection site in mice treated as indicated. Scale bars = 200 µm. (**C**) Quantification of hTau^+^/GFP^−^ cells per mm^2^ in mice treated as indicated. (**D)** Quantification of GFP^+^ cells per mm^2^ in mice treated as indicated. (**E**) Quantification of the hTau/GFP colocalization ratio in each condition, normalized to E3 CON condition. (**F**) Quantification of Tau spreading distance (μm) for each condition (*P-* values indicated on graphs; data presented as mean ± SD; one-way ANOVA with Tukey’s multiple comparisons test; n=5 mice/group). Each point represents an individual mouse.

## Discussion

Whether and how *APOE4*, the most common genetic Alzheimer’s disease risk factor, confers vulnerability to chronic stress and contributes to stress-related brain pathology has been unclear. In the current study, we find that GC-induced mitochondrial damage and ROS production, as well as tau hyperphosphorylation, oligomerization, and spreading, is exacerbated in *APOE4* humanized knockin mice compared to their *APOE3* counterparts, consistent with the reported vulnerability of *APOE4* carriers to stress^10,14,20,53-55^. Moreover, E4 animals exhibit mitochondrial dysfunction and enhanced GR activation at baseline, indicating that these underlying factors promote tau pathology and Alzheimer’s progression in the presence and absence of external stressors. Mechanistically, we show that the mPTP is a key driver of mitochondrial damage and tau pathology in the E4 background, and that mPTP inhibition is protective against both features in aging E4 mice. These findings shed light on the mechanisms of *APOE4*-mediated stress vulnerability and Alzheimer’s pathophysiology.

Multiple studies show a link between *APOE4* and mitochondrial dysfunction^18-21,35,39-41^, but the mechanisms by which ApoE4 causes this dysfunction remain unclear. In addition to its role as a secreted protein, ApoE localizes to mitochondria and mitochondria-associated endoplasmic reticulum membranes (MAMs) and is reported to interact with a number of mitochondrial and MAM proteins^41,56^, suggesting a role in these compartments. Moreover, C-terminally truncated ApoE4 fragments, the product of ApoE proteolytic cleavage in the neuronal cytosol^57^, were found to bind and inhibit the activity of mitochondrial respiratory chain proteins, leading to mitochondrial dysfunction^58^. These proteolytic fragments can also induce MAM formation and mitochondrial calcium overload, promote mitochondrial fission and fragmentation, and induce transcriptional changes in the nucleus^39,58,59^, all of which may contribute to mitochondrial dysfunction in neurons. In cardiomyocytes, ApoE bound to a specific form of low density lipoprotein was also observed to colocalize with the voltage-dependent anion-selective channel 1 (VDAC1) on the outer mitochondrial membrane and induce mPTP opening^60^, indicating a role for ApoE in regulating mPTP opening via VDAC1. Additional research will be required to determine whether this function is specific to, or enhanced by, the E4 variant.

In our previous study conducted in wild-type mice, we showed that mitochondrial damage precipitates tau pathology^17^, and we see an augmented version of this phenomenon in *APOE4* animals. Precisely how mitochondrial damage drives tau pathogenesis is unknown, but several mechanisms have been implicated. These include mROS-mediated oxidation of specific residues on tau (e.g., methionines), leading to tau cleavage and aggregation^61^, and leakage of a mitochondrial succinylating enzyme into the cytoplasm, inducing the succinylation and subsequent aggregation of tau^62^. There is also increasing evidence that ATP functions as a biological hydrotrope to maintain protein solubility in the cytosol, and that reduced levels of cytosolic ATP (e.g., resulting from mitochondrial dysfunction) contribute to the aggregation of intrinsically-disordered proteins such as tau^63,64^. All of these mechanisms could play a role in tau aggregation specifically, and more generally in the loss of proteostasis during chronic stress, aging, and Alzheimer’s pathogenesis, and thus merit future investigation.

An unexpected finding from this study is that GR activation is enhanced in the *APOE4* background. Several factors impact GR activation, including its chaperones, phosphorylation, and GC bioavailability^42^. Here, we show increased levels of the major GR co-chaperones Hsp90, Hsp70, and FKBP51 in brain tissue from E4 mice (Fig. **3**). This finding is consistent with another study showing upregulation of HSP70 and HSP90 transcripts in TE4 mice and their downregulation following neuron-specific *APOE4* deletion^23^. It was recently reported that hormone binding to GR is accelerated by Hsp90^43^, indicating a potential mechanism by which increased Hsp90 levels could enhance GR activation in E4 neurons during GC exposure. However, other mechanisms are likely responsible for the observed activation of GR in brains of E4 animals in the absence of exogenous/elevated GC levels (Fig. **3**). For instance, ApoE4 may alter the levels of enzymes involved in cortisol (for humans)/corticosterone (for mice) bioavailability. These include 11β-hydroxysteroid dehydrogenase, which converts inactive cortisone (or 11-dehydrocorticosterone, in the case of mice) to cortisol/corticosterone, and multidrug resistance P-glycoprotein (MDR1), a transporter that pumps GCs and other compounds out of cells^42^. Additionally, ApoE4 could up- or downregulate the levels of kinases and/or phosphatases that mediate GR phosphorylation (e.g., CDK2, JNK, p38 MAPK, PP5^42^), leading to higher basal GR activation. These possibilities will be tested in our ongoing studies.

Given the lack of effective Alzheimer’s disease therapies for *APOE4* carriers, who are highly susceptible to the most serious side effects of amyloid-clearing immunotherapy (i.e., amyloid-related imaging abnormalities with edema or hemorrhages, referred to as ARIA-E or ARIA-H)^65^, an important finding from our work is that inhibition of the mPTP is protective against ApoE4-mediated tau pathogenesis. Indeed, we show that mAPO prevents tau pathology when given for 2.5 months to young (∼1.5-month-old) TE4 mice prior to the onset of tauopathy-related phenotypes, and when given for a more limited, 15-day period to aging (15-16-month-old) E4 mice. In the young TE4 animals, mAPO administration not only protects against mitochondrial dysfunction and tau hyperphosphorylation/oligomerization in the hippocampus, but also prevents synapse loss, indicating amelioration of the synaptotoxic effects of pathogenic tau associated with cognitive impairment. Similarly, mAPO administration to aging E4 animals normalizes their mitochondrial function, tau phosphorylation/oligomerization, and tau spreading to the levels of E3 control animals, suggesting that mPTP inhibition in middle age can effectively counteract ApoE4-driven pathology. While our experiments here have focused on mitochondrial dysfunction and tau pathology, the other major Alzheimer’s pathomechanisms, amyloid-beta (Aβ) production and neuroinflammation, are also augmented by stress/GCs and ApoE4, and recent studies suggest that they would also be mitigated by mPTP inhibition. Indeed, Aβ was shown to interact with the mitochondrial membrane and trigger opening of the mPTP, while CypD inhibitors or knockout have protective effects against Aβ-induced toxicity in animal models^66,67^. In addition, mPTP opening is reported to trigger activation of pro-inflammatory pathways through the release of mitochondrial DNA into the cytosol^68-70^, and CypD inhibitors or knockout reduce pro-inflammatory signaling in animal models of acute inflammation and Alzheimer’s disease^67,71,72^. Other studies show that mAPO administration or CypD inhibition/knockout are protective against neurodegeneration in animal models of Alzheimer’s and Parkinson’s diseases and multiple sclerosis^66,73-78^, demonstrating the broader therapeutic promise of targeting the mPTP. Although the specific CypD inhibitor CsA doesn’t cross the blood-brain-barrier and thus cannot be readily used to target the central nervous system^79^, other inhibitors are currently under development^67,80^ and represent an exciting new category of drugs for the treatment of neurodegenerative disease.

In summary, we find that mitochondrial dysfunction underlies chronic stress- and ApoE4-induced tau phosphorylation, oligomerization, and spreading, and that inhibition of the mPTP prevents these effects in *APOE4* carriers. This strategy may represent a viable therapeutic option for preventing stress- and ApoE4-mediated neurodegeneration in Alzheimer’s disease.

## Abbreviations

AAV: adeno-associated virus
Aβ: amyloid-beta
APOE: apolipoprotein E
ARIA-E/ARIA-H: amyloid-related imaging abnormalities with edema or hemorrhages
CDK2: cyclin-dependent kinase 2
CsA: cyclosporin A
CypD: cyclophilin D
DEX: dexamethasone
E3/E4 mice: *APOE3/APOE4* targeted replacement mice
GC: glucocorticoids
GFP: green fluorescent protein
GR: glucocorticoid receptor
Hsp 70/90: heat shock protein 70/90
hTau: human tau
JNK: c-Jun N-terminal kinase
MAM: mitochondria-associated endoplasmic reticulum membrane
mAPO: mito-apocynin
mPTP: mitochondrial permeability transition pore
mROS: mitochondrial reactive oxygen species
NADPH: nicotinamide adenine dinucleotide phosphate
p38 MAPK: mitogen-activated protein kinase
PP5: protein phosphatase 5
TE3/TE4 mice: PS19 tauopathy mice crossed with *APOE3/APOE4* targeted replacement mice
VDAC1: voltage-dependent anion-selective channel 1

## Acknowledgements

We would like to thank members of the Laboratory of Mitochondrial Function, Institute for Basic Research in Developmental Disabilities, for use of the EPR, and Ioannis Sotiropoulos and members of the Sotiropoulos lab for many helpful discussions about stress-related brain pathology.

## Supplementary Material

Supplementary material is available at *Brain* online.

## Author Contributions

Q.Y., F.D., and C.L.W. designed the research; Q.Y., F.D., and J.G. performed experiments and analyzed data; C.L.W. wrote the manuscript, Q.Y. and F.D. prepared figures and edited the manuscript. All authors read and approved the final manuscript.

## References

1. Serrano-Pozo, A., Das, S., and Hyman, B.T. (2021). APOE and Alzheimer’s disease: advances in genetics, pathophysiology, and therapeutic approaches. Lancet Neurol 20, 68–80. 10.1016/S1474-4422(20)30412-9.

2. Mravec, B., Horvathova, L., and Padova, A. (2018). Brain Under Stress and Alzheimer’s Disease. Cell Mol Neurobiol 38, 73–84. 10.1007/s10571-017-0521-1.

3. Caruso, A., Nicoletti, F., Gaetano, A., and Scaccianoce, S. (2019). Risk Factors for Alzheimer’s Disease: Focus on Stress. Front Pharmacol 10, 976. 10.3389/fphar.2019.00976.

4. Burke, M.R., Sotiropoulos, I., and Waites, C.L. (2024). The multiple roles of chronic stress and glucocorticoids in Alzheimer’s disease pathogenesis. Trends Neurosci. 10.1016/j.tins.2024.08.015.

5. Machado, A., Herrera, A.J., de Pablos, R.M., Espinosa-Oliva, A.M., Sarmiento, M., Ayala, A., Venero, J.L., Santiago, M., Villaran, R.F., Delgado-Cortes, M.J., et al. (2014). Chronic stress as a risk factor for Alzheimer’s disease. Rev Neurosci 25, 785–804. 10.1515/revneuro-2014-0035.

6. Johansson, L., Guo, X., Waern, M., Ostling, S., Gustafson, D., Bengtsson, C., and Skoog, I. (2010). Midlife psychological stress and risk of dementia: a 35-year longitudinal population study. Brain 133, 2217–2224. 10.1093/brain/awq116.

7. Huang, C.W., Lui, C.C., Chang, W.N., Lu, C.H., Wang, Y.L., and Chang, C.C. (2009). Elevated basal cortisol level predicts lower hippocampal volume and cognitive decline in Alzheimer’s disease. J Clin Neurosci 16, 1283–1286. 10.1016/j.jocn.2008.12.026.

8. Holmquist, S., Nordstrom, A., and Nordstrom, P. (2020). The association of depression with subsequent dementia diagnosis: A Swedish nationwide cohort study from 1964 to 2016. PLoS Med 17, e1003016. 10.1371/journal.pmed.1003016.

9. Saiz-Vazquez, O., Gracia-Garcia, P., Ubillos-Landa, S., Puente-Martinez, A., Casado-Yusta, S., Olaya, B., and Santabarbara, J. (2021). Depression as a Risk Factor for Alzheimer’s Disease: A Systematic Review of Longitudinal Meta-Analyses. J Clin Med 10. 10.3390/jcm10091809.

10. Yen, Y.C., Rebok, G.W., Yang, M.J., and Lung, F.W. (2008). A multilevel analysis of the influence of Apolipoprotein E genotypes on depressive symptoms in late-life moderated by the environment. Prog Neuropsychopharmacol Biol Psychiatry 32, 479–486. 10.1016/j.pnpbp.2007.09.023.

11. Yen, Y.C., Rebok, G.W., Gallo, J.J., Yang, M.J., Lung, F.W., and Shih, C.H. (2007). ApoE4 allele is associated with late-life depression: a population-based study. The American journal of geriatric psychiatry : official journal of the American Association for Geriatric Psychiatry 15, 858–868. 10.1097/JGP.0b013e3180f63373.

12. Holmes, S.E., Esterlis, I., Mazure, C.M., Lim, Y.Y., Ames, D., Rainey-Smith, S., Martins, R.N., Salvado, O., Dore, V., Villemagne, V.L., et al. (2016). beta-Amyloid, APOE and BDNF Genotype, and Depressive and Anxiety Symptoms in Cognitively Normal Older Women and Men. The American journal of geriatric psychiatry : official journal of the American Association for Geriatric Psychiatry 24, 1191–1195. 10.1016/j.jagp.2016.08.007.

13. Dose, J., Huebbe, P., Nebel, A., and Rimbach, G. (2016). APOE genotype and stress response - a mini review. Lipids Health Dis 15, 121. 10.1186/s12944-016-0288-2.

14. Peavy, G.M., Lange, K.L., Salmon, D.P., Patterson, T.L., Goldman, S., Gamst, A.C., Mills, P.J., Khandrika, S., and Galasko, D. (2007). The effects of prolonged stress and APOE genotype on memory and cortisol in older adults. Biol Psychiatry 62, 472–478. 10.1016/j.biopsych.2007.03.013.

15. Sotiropoulos, I., Silva, J.M., Gomes, P., Sousa, N., and Almeida, O.F.X. (2019). Stress and the Etiopathogenesis of Alzheimer’s Disease and Depression. Adv Exp Med Biol 1184, 241–257. 10.1007/978-981-32-9358-8_20.

16. Wang, W., Zhao, F., Ma, X., Perry, G., and Zhu, X. (2020). Mitochondria dysfunction in the pathogenesis of Alzheimer’s disease: recent advances. Molecular neurodegeneration 15, 30. 10.1186/s13024-020-00376-6.

17. Du, F., Yu, Q., Swerdlow, R.H., and Waites, C.L. (2023). Glucocorticoid-driven mitochondrial damage stimulates Tau pathology. Brain. 10.1093/brain/awad127.

18. Yin, J., Reiman, E.M., Beach, T.G., Serrano, G.E., Sabbagh, M.N., Nielsen, M., Caselli, R.J., and Shi, J. (2020). Effect of ApoE isoforms on mitochondria in Alzheimer disease. Neurology 94, e2404–e2411. 10.1212/WNL.0000000000009582.

19. Simonovitch, S., Schmukler, E., Masliah, E., Pinkas-Kramarski, R., and Michaelson, D.M. (2019). The Effects of APOE4 on Mitochondrial Dynamics and Proteins in vivo. J Alzheimers Dis 70, 861–875. 10.3233/JAD-190074.

20. Fang, W., Xiao, N., Zeng, G., Bi, D., Dai, X., Mi, X., Ye, Q., Chen, X., and Zhang, J. (2021). APOE4 genotype exacerbates the depression-like behavior of mice during aging through ATP decline. Transl Psychiatry 11, 507. 10.1038/s41398-021-01631-0.

21. Schmukler, E., Solomon, S., Simonovitch, S., Goldshmit, Y., Wolfson, E., Michaelson, D.M., and Pinkas-Kramarski, R. (2020). Altered mitochondrial dynamics and function in APOE4-expressing astrocytes. Cell Death Dis 11, 578. 10.1038/s41419-020-02776-4.

22. Shi, Y., Yamada, K., Liddelow, S.A., Smith, S.T., Zhao, L., Luo, W., Tsai, R.M., Spina, S., Grinberg, L.T., Rojas, J.C., et al. (2017). ApoE4 markedly exacerbates tau-mediated neurodegeneration in a mouse model of tauopathy. Nature 549, 523–527. 10.1038/nature24016.

23. Koutsodendris, N., Blumenfeld, J., Agrawal, A., Traglia, M., Grone, B., Zilberter, M., Yip, O., Rao, A., Nelson, M.R., Hao, Y., et al. (2023). Neuronal APOE4 removal protects against tau-mediated gliosis, neurodegeneration and myelin deficits. Nat Aging 3, 275–296. 10.1038/s43587-023-00368-3.

24. Wang, C., Xiong, M., Gratuze, M., Bao, X., Shi, Y., Andhey, P.S., Manis, M., Schroeder, C., Yin, Z., Madore, C., et al. (2021). Selective removal of astrocytic APOE4 strongly protects against tau-mediated neurodegeneration and decreases synaptic phagocytosis by microglia. Neuron 109, 1657–1674 e1657. 10.1016/j.neuron.2021.03.024.

25. Du, F., Yu, Q., Swerdlow, R.H., and Waites, C.L. (2023). Glucocorticoid-driven mitochondrial damage stimulates Tau pathology. Brain 146, 4378–4394. 10.1093/brain/awad127.

26. Du, F., Yu, Q., Yan, S., Hu, G., Lue, L.F., Walker, D.G., Wu, L., Yan, S.F., Tieu, K., and Yan, S.S. (2017). PINK1 signalling rescues amyloid pathology and mitochondrial dysfunction in Alzheimer’s disease. Brain 140, 3233–3251. 10.1093/brain/awx258.

27. Yu, Q., Du, F., Belli, I., Gomes, P.A., Sotiropoulos, I., and Waites, C.L. (2024). Glucocorticoid stress hormones stimulate vesicle-free Tau secretion and spreading in the brain. Cell Death Dis 15, 73. 10.1038/s41419-024-06458-3.

28. Fang, D., Zhang, Z., Li, H., Yu, Q., Douglas, J.T., Bratasz, A., Kuppusamy, P., and Yan, S.S. (2016). Increased Electron Paramagnetic Resonance Signal Correlates with Mitochondrial Dysfunction and Oxidative Stress in an Alzheimer’s disease Mouse Brain. J Alzheimers Dis 51, 571–580. 10.3233/JAD-150917.

29. Rauch, J.N., Luna, G., Guzman, E., Audouard, M., Challis, C., Sibih, Y.E., Leshuk, C., Hernandez, I., Wegmann, S., Hyman, B.T., et al. (2020). LRP1 is a master regulator of tau uptake and spread. Nature 580, 381–385. 10.1038/s41586-020-2156-5.

30. Wegmann, S., Bennett, R.E., Delorme, L., Robbins, A.B., Hu, M., McKenzie, D., Kirk, M.J., Schiantarelli, J., Tunio, N., Amaral, A.C., et al. (2019). Experimental evidence for the age dependence of tau protein spread in the brain. Sci Adv 5, eaaw6404. 10.1126/sciadv.aaw6404.

31. You, J.M., Yun, S.J., Nam, K.N., Kang, C., Won, R., and Lee, E.H. (2009). Mechanism of glucocorticoid-induced oxidative stress in rat hippocampal slice cultures. Can J Physiol Pharmacol 87, 440–447. 10.1139/y09-027.

32. Vaz-Silva, J., Gomes, P., Jin, Q., Zhu, M., Zhuravleva, V., Quintremil, S., Meira, T., Silva, J., Dioli, C., Soares-Cunha, C., et al. (2018). Endolysosomal degradation of Tau and its role in glucocorticoid-driven hippocampal malfunction. EMBO J. 10.15252/embj.201899084.

33. Yang, L., Zhou, H., Huang, L., Su, Y., Kong, L., Ji, P., Sun, R., Wang, C., Li, W., and Li, W. (2022). Stress level of glucocorticoid exacerbates neuronal damage and Abeta production through activating NLRP1 inflammasome in primary cultured hippocampal neurons of APP-PS1 mice. Int Immunopharmacol 110, 108972. 10.1016/j.intimp.2022.108972.

34. Sotiropoulos, I., Catania, C., Riedemann, T., Fry, J.P., Breen, K.C., Michaelidis, T.M., and Almeida, O.F. (2008). Glucocorticoids trigger Alzheimer disease-like pathobiochemistry in rat neuronal cells expressing human tau. J Neurochem 107, 385–397. 10.1111/j.1471-4159.2008.05613.x.

35. Area-Gomez, E., Larrea, D., Pera, M., Agrawal, R.R., Guilfoyle, D.N., Pirhaji, L., Shannon, K., Arain, H.A., Ashok, A., Chen, Q., et al. (2020). APOE4 is Associated with Differential Regional Vulnerability to Bioenergetic Deficits in Aged APOE Mice. Scientific reports 10, 4277. 10.1038/s41598-020-61142-8.

36. Merezhko, M., Brunello, C.A., Yan, X., Vihinen, H., Jokitalo, E., Uronen, R.L., and Huttunen, H.J. (2018). Secretion of Tau via an Unconventional Non-vesicular Mechanism. Cell reports 25, 2027–2035 e2024. 10.1016/j.celrep.2018.10.078.

37. Katsinelos, T., Zeitler, M., Dimou, E., Karakatsani, A., Muller, H.M., Nachman, E., Steringer, J.P., Ruiz de Almodovar, C., Nickel, W., and Jahn, T.R. (2018). Unconventional Secretion Mediates the Trans-cellular Spreading of Tau. Cell reports 23, 2039–2055. 10.1016/j.celrep.2018.04.056.

38. Yu, Q., Du, F., Belli, I., Gomes, P.A., Sotiropoulos, I., and Waites, C.L. (2023). Glucocorticoid stress hormones stimulate vesicle-free Tau secretion and spreading in the brain. bioRxiv. 10.1101/2023.06.07.544054.

39. Orr, A.L., Kim, C., Jimenez-Morales, D., Newton, B.W., Johnson, J.R., Krogan, N.J., Swaney, D.L., and Mahley, R.W. (2019). Neuronal Apolipoprotein E4 Expression Results in Proteome-Wide Alterations and Compromises Bioenergetic Capacity by Disrupting Mitochondrial Function. J Alzheimers Dis 68, 991–1011. 10.3233/JAD-181184.

40. Qi, G., Mi, Y., Shi, X., Gu, H., Brinton, R.D., and Yin, F. (2021). ApoE4 Impairs Neuron-Astrocyte Coupling of Fatty Acid Metabolism. Cell reports 34, 108572. 10.1016/j.celrep.2020.108572.

41. Mahley, R.W. (2023). Apolipoprotein E4 targets mitochondria and the mitochondria-associated membrane complex in neuropathology, including Alzheimer’s disease. Curr Opin Neurobiol 79, 102684. 10.1016/j.conb.2023.102684.

42. Oakley, R.H., and Cidlowski, J.A. (2013). The biology of the glucocorticoid receptor: new signaling mechanisms in health and disease. J Allergy Clin Immunol 132, 1033–1044. 10.1016/j.jaci.2013.09.007.

43. Kaziales, A., Barkovits, K., Marcus, K., and Richter, K. (2020). Glucocorticoid receptor complexes form cooperatively with the Hsp90 co-chaperones Pp5 and FKBPs. Scientific reports 10, 10733. 10.1038/s41598-020-67645-8.

44. Kirschke, E., Goswami, D., Southworth, D., Griffin, P.R., and Agard, D.A. (2014). Glucocorticoid receptor function regulated by coordinated action of the Hsp90 and Hsp70 chaperone cycles. Cell 157, 1685–1697. 10.1016/j.cell.2014.04.038.

45. Bonora, M., Giorgi, C., and Pinton, P. (2022). Molecular mechanisms and consequences of mitochondrial permeability transition. Nat Rev Mol Cell Biol 23, 266–285. 10.1038/s41580-021-00433-y.

46. Bonora, M., Morganti, C., Morciano, G., Giorgi, C., Wieckowski, M.R., and Pinton, P. (2016). Comprehensive analysis of mitochondrial permeability transition pore activity in living cells using fluorescence-imaging-based techniques. Nat Protoc 11, 1067–1080. 10.1038/nprot.2016.064.

47. Parhizkar, S., and Holtzman, D.M. (2022). APOE mediated neuroinflammation and neurodegeneration in Alzheimer’s disease. Semin Immunol 59, 101594. 10.1016/j.smim.2022.101594.

48. Koriath, C., Lashley, T., Taylor, W., Druyeh, R., Dimitriadis, A., Denning, N., Williams, J., Warren, J.D., Fox, N.C., Schott, J.M., et al. (2019). ApoE4 lowers age at onset in patients with frontotemporal dementia and tauopathy independent of amyloid-beta copathology. Alzheimers Dement (Amst) 11, 277–280. 10.1016/j.dadm.2019.01.010.

49. Wu, M., Zhang, M., Yin, X., Chen, K., Hu, Z., Zhou, Q., Cao, X., Chen, Z., and Liu, D. (2021). The role of pathological tau in synaptic dysfunction in Alzheimer’s diseases. Translational neurodegeneration 10, 45. 10.1186/s40035-021-00270-1.

50. Vyas, S., Rodrigues, A.J., Silva, J.M., Tronche, F., Almeida, O.F., Sousa, N., and Sotiropoulos, I. (2016). Chronic Stress and Glucocorticoids: From Neuronal Plasticity to Neurodegeneration. Neural plasticity 2016, 6391686. 10.1155/2016/6391686.

51. Steward, A., Biel, D., Dewenter, A., Roemer, S., Wagner, F., Dehsarvi, A., Rathore, S., Otero Svaldi, D., Higgins, I., Brendel, M., et al. (2023). ApoE4 and Connectivity-Mediated Spreading of Tau Pathology at Lower Amyloid Levels. JAMA Neurol 80, 1295–1306. 10.1001/jamaneurol.2023.4038.

52. Saroja, S.R., Gorbachev, K., Julia, T., Goate, A.M., and Pereira, A.C. (2022). Astrocyte-secreted glypican-4 drives APOE4-dependent tau hyperphosphorylation. Proc Natl Acad Sci U S A 119, e2108870119. 10.1073/pnas.2108870119.

53. Lin, L.Y., Zhang, J., Dai, X.M., Xiao, N.A., Wu, X.L., Wei, Z., Fang, W.T., Zhu, Y.G., and Chen, X.C. (2016). Early-life stress leads to impaired spatial learning and memory in middle-aged ApoE4-TR mice. Molecular neurodegeneration 11, 51. 10.1186/s13024-016-0107-2.

54. Zeng, Y., Hughes, C.L., Lewis, M.A., Li, J., and Zhang, F. (2011). Interactions between life stress factors and carrying the APOE4 allele adversely impact self-reported health in old adults. J Gerontol A Biol Sci Med Sci 66, 1054–1061. 10.1093/gerona/glr106.

55. Zhang, J., Lin, L., Dai, X., Xiao, N., Ye, Q., and Chen, X. (2021). ApoE4 increases susceptibility to stress-induced age-dependent depression-like behavior and cognitive impairment. J Psychiatr Res 143, 292–301. 10.1016/j.jpsychires.2021.09.029.

56. Rueter, J., Rimbach, G., and Huebbe, P. (2022). Functional diversity of apolipoprotein E: from subcellular localization to mitochondrial function. Cell Mol Life Sci 79, 499. 10.1007/s00018-022-04516-7.

57. Munoz, S.S., Garner, B., and Ooi, L. (2019). Understanding the Role of ApoE Fragments in Alzheimer’s Disease. Neurochem Res 44, 1297–1305. 10.1007/s11064-018-2629-1.

58. Nakamura, T., Watanabe, A., Fujino, T., Hosono, T., and Michikawa, M. (2009). Apolipoprotein E4 (1-272) fragment is associated with mitochondrial proteins and affects mitochondrial function in neuronal cells. Molecular neurodegeneration 4, 35. 10.1186/1750-1326-4-35.

59. Liang, T., Hang, W., Chen, J., Wu, Y., Wen, B., Xu, K., Ding, B., and Chen, J. (2021). ApoE4 (Delta272-299) induces mitochondrial-associated membrane formation and mitochondrial impairment by enhancing GRP75-modulated mitochondrial calcium overload in neuron. Cell Biosci 11, 50. 10.1186/s13578-021-00563-y.

60. Chen, W.Y., Chen, Y.F., Chan, H.C., Chung, C.H., Peng, H.Y., Ho, Y.C., Chen, C.H., Chang, K.C., Tang, C.H., and Lee, A.S. (2020). Role of apolipoprotein E in electronegative low-density lipoprotein-induced mitochondrial dysfunction in cardiomyocytes. Metabolism 107, 154227. 10.1016/j.metabol.2020.154227.

61. Samelson, A.J., Ariqat, N., McKetney, J., Rohanitazangi, G., Bravo, C.P., Bose, R., Travaglini, K.J., Lam, V.L., Goodness, D., Dixon, G., et al. (2024). CRISPR screens in iPSC-derived neurons reveal principles of tau proteostasis. bioRxiv. 10.1101/2023.06.16.545386.

62. Yang, Y., Tapias, V., Acosta, D., Xu, H., Chen, H., Bhawal, R., Anderson, E.T., Ivanova, E., Lin, H., Sagdullaev, B.T., et al. (2022). Altered succinylation of mitochondrial proteins, APP and tau in Alzheimer’s disease. Nature communications 13, 159. 10.1038/s41467-021-27572-2.

63. Patel, A., Malinovska, L., Saha, S., Wang, J., Alberti, S., Krishnan, Y., and Hyman, A.A. (2017). ATP as a biological hydrotrope. Science 356, 753–756. 10.1126/science.aaf6846.

64. Sridharan, S., Kurzawa, N., Werner, T., Gunthner, I., Helm, D., Huber, W., Bantscheff, M., and Savitski, M.M. (2019). Proteome-wide solubility and thermal stability profiling reveals distinct regulatory roles for ATP. Nature communications 10, 1155. 10.1038/s41467-019-09107-y.

65. Sato, K., Niimi, Y., Ihara, R., Suzuki, K., Iwata, A., and Iwatsubo, T. (2023). APOE-epsilon4 allele[s]-associated adverse events reported from placebo arm in clinical trials for Alzheimer’s disease: implications for anti-amyloid beta therapy. Front Dement 2, 1320329. 10.3389/frdem.2023.1320329.

66. Du, H., Guo, L., Fang, F., Chen, D., Sosunov, A.A., McKhann, G.M., Yan, Y., Wang, C., Zhang, H., Molkentin, J.D., et al. (2008). Cyclophilin D deficiency attenuates mitochondrial and neuronal perturbation and ameliorates learning and memory in Alzheimer’s disease. Nat Med 14, 1097–1105. 10.1038/nm.1868.

67. Samanta, S., Akhter, F., Roy, A., Chen, D., Turner, B., Wang, Y., Clemente, N., Wang, C., Swerdlow, R.H., Battaile, K.P., et al. (2023). New cyclophilin D inhibitor rescues mitochondrial and cognitive function in Alzheimer’s disease. Brain. 10.1093/brain/awad432.

68. Yu, C.H., Davidson, S., Harapas, C.R., Hilton, J.B., Mlodzianoski, M.J., Laohamonthonkul, P., Louis, C., Low, R.R.J., Moecking, J., De Nardo, D., et al. (2020). TDP-43 Triggers Mitochondrial DNA Release via mPTP to Activate cGAS/STING in ALS. Cell 183, 636–649 e618. 10.1016/j.cell.2020.09.020.

69. Xian, H., Watari, K., Sanchez-Lopez, E., Offenberger, J., Onyuru, J., Sampath, H., Ying, W., Hoffman, H.M., Shadel, G.S., and Karin, M. (2022). Oxidized DNA fragments exit mitochondria via mPTP- and VDAC-dependent channels to activate NLRP3 inflammasome and interferon signaling. Immunity 55, 1370–1385 e1378. 10.1016/j.immuni.2022.06.007.

70. Marchi, S., Guilbaud, E., Tait, S.W.G., Yamazaki, T., and Galluzzi, L. (2023). Mitochondrial control of inflammation. Nat Rev Immunol 23, 159–173. 10.1038/s41577-022-00760-x.

71. Gui, Q., Jiang, Z., and Zhang, L. (2021). Insights into the modulatory role of cyclosporine A and its research advances in acute inflammation. Int Immunopharmacol 93, 107420. 10.1016/j.intimp.2021.107420.

72. Guareschi, F., Fonseca, C., Silva, S., Pescina, S., Nicoli, S., Buttini, F., Sonvico, F., and Fortuna, A. (2024). Therapeutic effect of cyclosporine A-loading TPGS micelles on a mouse model of LPS-induced neuroinflammation. Eur J Pharm Sci 205, 106994. 10.1016/j.ejps.2024.106994.

73. Langley, M., Ghosh, A., Charli, A., Sarkar, S., Ay, M., Luo, J., Zielonka, J., Brenza, T., Bennett, B., Jin, H., et al. (2017). Mito-Apocynin Prevents Mitochondrial Dysfunction, Microglial Activation, Oxidative Damage, and Progressive Neurodegeneration in MitoPark Transgenic Mice. Antioxid Redox Signal 27, 1048–1066. 10.1089/ars.2016.6905.

74. Liu, N., Lin, M.M., Huang, S.S., Liu, Z.Q., Wu, J.C., Liang, Z.Q., Qin, Z.H., and Wang, Y. (2021). NADPH and Mito-Apocynin Treatment Protects Against KA-Induced Excitotoxic Injury Through Autophagy Pathway. Front Cell Dev Biol 9, 612554. 10.3389/fcell.2021.612554.

75. Ghosh, A., Langley, M.R., Harischandra, D.S., Neal, M.L., Jin, H., Anantharam, V., Joseph, J., Brenza, T., Narasimhan, B., Kanthasamy, A., et al. (2016). Mitoapocynin Treatment Protects Against Neuroinflammation and Dopaminergic Neurodegeneration in a Preclinical Animal Model of Parkinson’s Disease. J Neuroimmune Pharmacol 11, 259–278. 10.1007/s11481-016-9650-4.

76. Dranka, B.P., Gifford, A., McAllister, D., Zielonka, J., Joseph, J., O’Hara, C.L., Stucky, C.L., Kanthasamy, A.G., and Kalyanaraman, B. (2014). A novel mitochondrially-targeted apocynin derivative prevents hyposmia and loss of motor function in the leucine-rich repeat kinase 2 (LRRK2(R1441G)) transgenic mouse model of Parkinson’s disease. Neurosci Lett 583, 159–164. 10.1016/j.neulet.2014.09.042.

77. Du, H., Guo, L., Zhang, W., Rydzewska, M., and Yan, S. (2011). Cyclophilin D deficiency improves mitochondrial function and learning/memory in aging Alzheimer disease mouse model. Neurobiol Aging 32, 398–406. 10.1016/j.neurobiolaging.2009.03.003.

78. Warne, J., Pryce, G., Hill, J.M., Shi, X., Lenneras, F., Puentes, F., Kip, M., Hilditch, L., Walker, P., Simone, M.I., et al. (2016). Selective Inhibition of the Mitochondrial Permeability Transition Pore Protects against Neurodegeneration in Experimental Multiple Sclerosis. J Biol Chem 291, 4356–4373. 10.1074/jbc.M115.700385.

79. Bellwon, P., Culot, M., Wilmes, A., Schmidt, T., Zurich, M.G., Schultz, L., Schmal, O., Gramowski-Voss, A., Weiss, D.G., Jennings, P., et al. (2015). Cyclosporine A kinetics in brain cell cultures and its potential of crossing the blood-brain barrier. Toxicol In Vitro 30, 166–175. 10.1016/j.tiv.2015.01.003.

80. Haleckova, A., Benek, O., Zemanova, L., Dolezal, R., and Musilek, K. (2022). Small-molecule inhibitors of cyclophilin D as potential therapeutics in mitochondria-related diseases. Med Res Rev 42, 1822–1855. 10.1002/med.21892.

