## Supplemental figures and legends for "*APOE4* exacerbates glucocorticoid stress hormone-induced tau pathology via mitochondrial dysfunction"

**Supplemental Material**

**Supplemental Figure Legends**

**Figure S1: Dexamethasone suppresses endogenous blood corticosterone and induces body weight loss.** (**A**) Quantification of corticosterone levels measured by ELISA in serum from 9-10-month-old E3/E4 mice treated with vehicle (CON) or dexamethasone (DEX)(*P*-values indicated on graphs; unpaired *t*-test; n=5 mice per group). (**B**) Quantification of body weight loss in the indicated mice (**$P_{E3CON\ vs.\ E3DEX}$ <0.01 (days 13-15), **$P_{E4CON\ vs.\ E4DEX}$ <0.01 (days 13-15), #$P_{E3CON\ vs.\ E3DEX}$ <0.05 (days 11-12), #$P_{E4CON\ vs.\ E4DEX}$ <0.01 (days 10-12); two-way ANOVA with multiple comparisons; n=10 mice/condition). All data presented as mean ± SD.

**Figure S2: APOE4 carriers exhibit mitochondrial dysfunction and tau pathology that is rescued by inhibitors of mPTP opening.** (**A**-**B**) Complex I activity (**A**) and ATP levels (**B**) in E3/E4 hippocampal neurons treated with vehicle (CON) or dexamethasone (DEX), normalized to the E3 CON condition (*P*-values are indicated on the graphs; data presented as mean ± SD; one-way ANOVA with Tukey's multiple comparisons test; n=5 samples/condition). (**C**-**D**) Quantification of TOMA-1 (**C**) and MitoSOX (**D**) fluorescence intensity in primary 14 DIV E4 hippocampal neurons treated with vehicle (CON), DEX, DEX + mito-apocynin (mAPO), or DEX + cyclosporin A (CsA). Intensity values are normalized to the DEX condition (*P*-values indicated on graphs; data presented as mean ± SD; one-way ANOVA with Tukey's multiple comparisons test; n=8 fields of view/condition). (**E-F**) Complex I activity (**E**) and ATP levels (**F**) in hippocampal tissue of 3.5-4-month-old TE3/TE4 mice treated with vehicle (CON) or mito-apocynin (mAPO), normalized to the TE3 condition (*P*-values indicated on graphs; data presented as mean ± SD; one-way ANOVA with Tukey's multiple comparisons test; n=5-6 samples/condition). Each point represents an individual mouse.

**Figure S3: Full immunoblots for all figures.** Rectangular boxes indicate the regions shown in the indicated figures. Proteins are labeled on the left and sizes (kD) on the right.

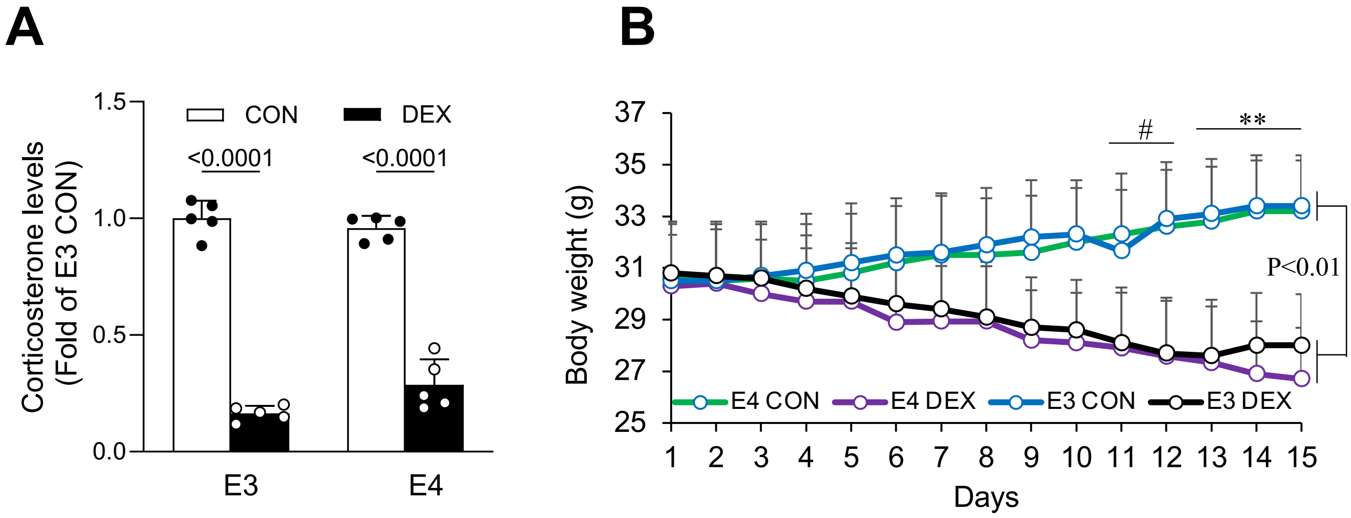
A
B
1.5
1.0
0.5
0.0
CON
DEX
<0.0001
<0.0001
Corticosterone levels
(Fold of E3 CON)
E3
E4
37
35
33
31
29
27
25
Body weight (g)
#
**
P<0.01
E4 CON
E4 DEX
E3 CON
E3 DEX
1 2 3 4 5 6 7 8 9 10 11 12 13 14 15
Days

Fig. S1

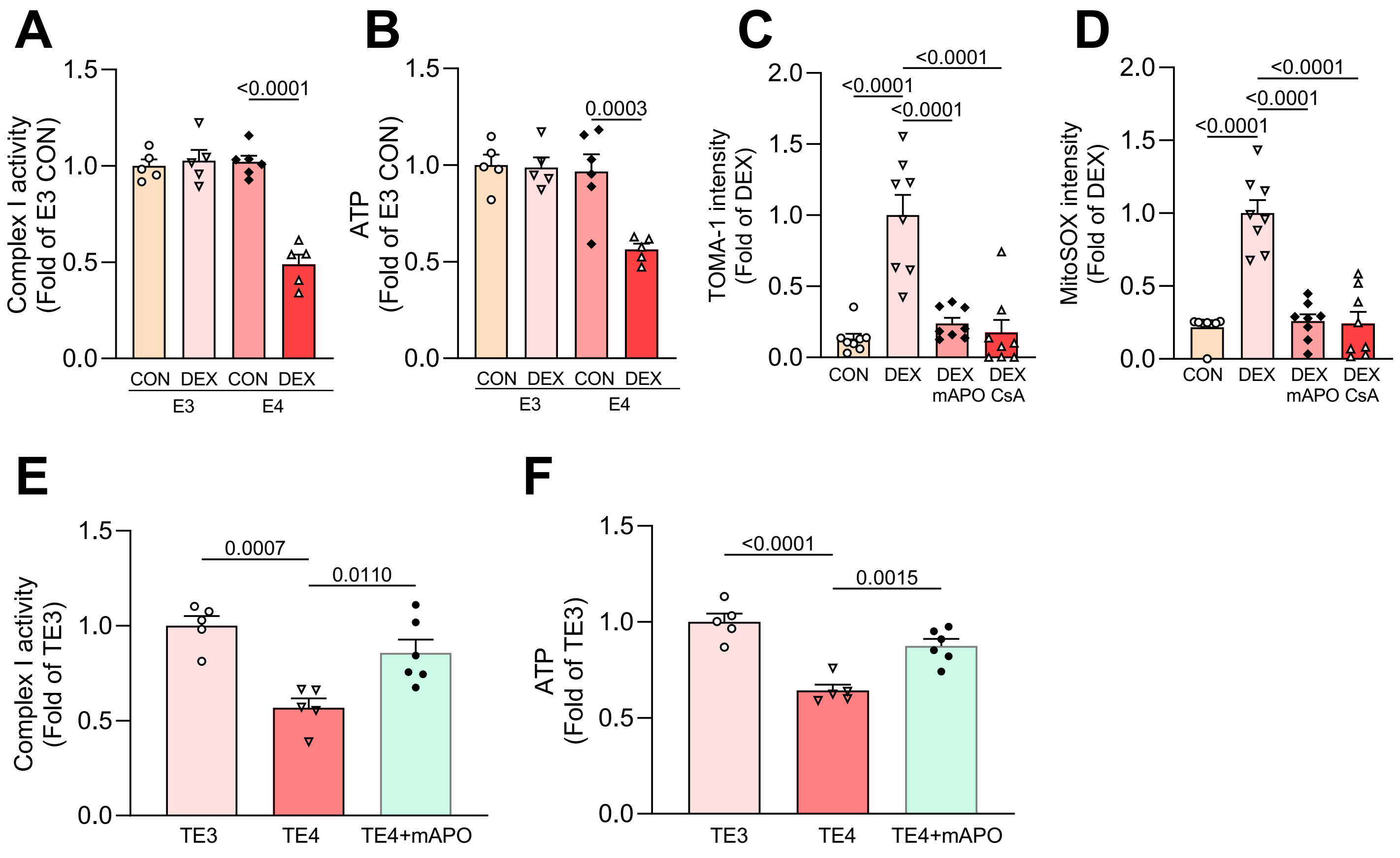
A
B
C
D
E
F
Complex I activity (Fold of E3 CON)
ATP (Fold of E3 CON)
TOMA-1 intensity (Fold of DEX)
MitoSOX intensity (Fold of DEX)
Complex I activity (Fold of TE3)
ATP (Fold of TE3)
<0.0001
0.0003
0.0007
0.0110
0.0015
CON DEX CON DEX
E3 E4
CON DEX DEX mAPO DEX CsA
TE3 TE4 TE4+mAPO

Fig. S2

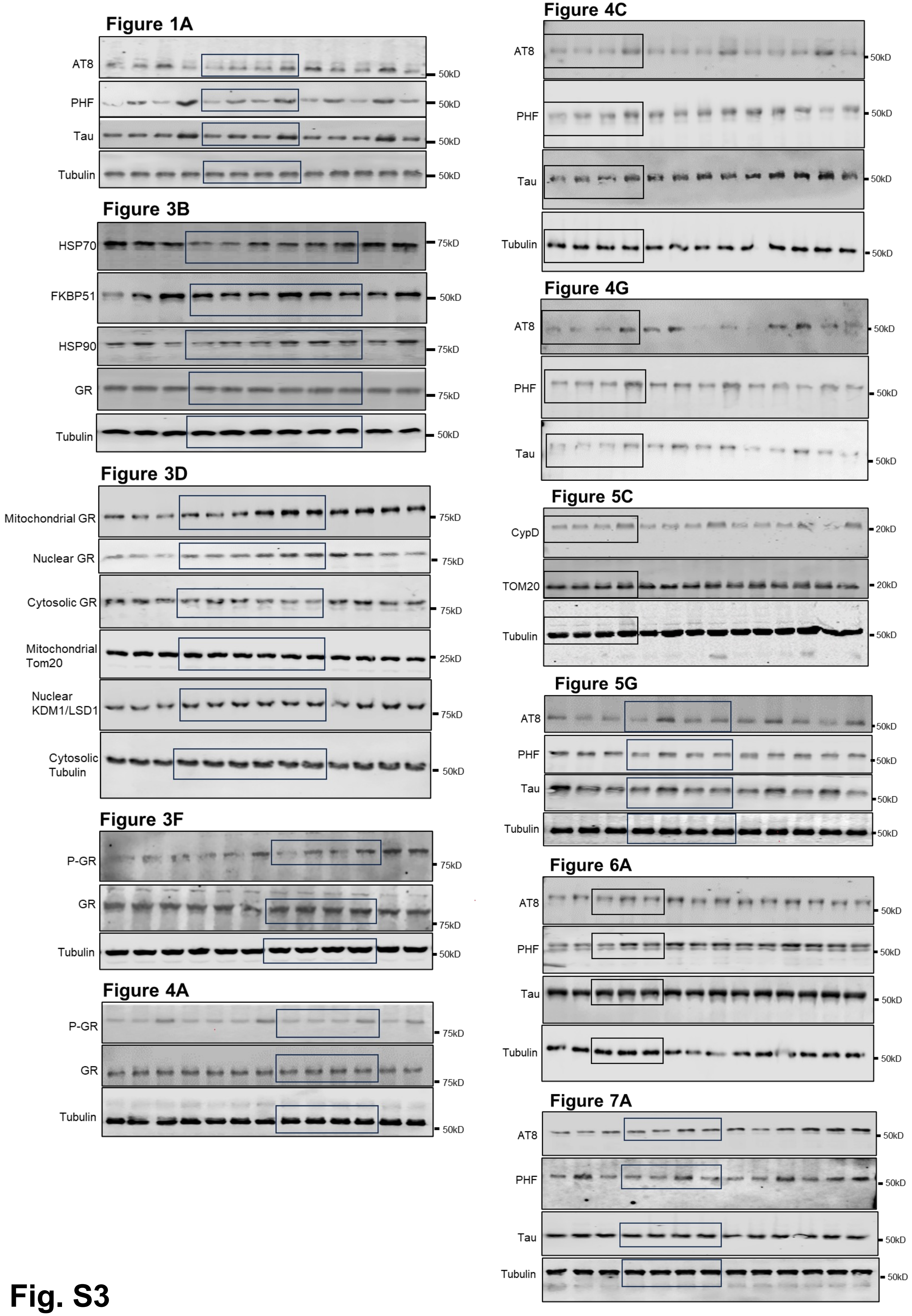

Figure 1A
AT8
PHF
Tau
Tubulin
50kD
Figure 3B
HSP70
FKBP51
HSP90
GR
Tubulin
75kD
50kD
Figure 3D
Mitochondrial GR
Nuclear GR
Cytosolic GR
Mitochondrial Tom20
Nuclear KDM1/LSD1
Cytosolic Tubulin
75kD
25kD
50kD
Figure 3F
P-GR
GR
Tubulin
Figure 4A
P-GR
GR
Tubulin
Figure 4C
AT8
PHF
Tau
Tubulin
Figure 4G
AT8
PHF
Tau
Figure 5C
CypD
TOM20
Tubulin
20kD
Figure 5G
AT8
PHF
Tau
Tubulin
Figure 6A
AT8
PHF
Tau
Tubulin
Figure 7A
AT8
PHF
Tau
Tubulin


Fig. S3
